# Multi-site phosphorylation of BCL2L13 in a TBK1- and AMPK-dependent manner reveals new modes of mitophagy regulation

**DOI:** 10.1101/2025.07.10.663832

**Authors:** Hoda Ahmed, Mack B. Reynolds, Gary Kasof, Gerald S. Shadel, Reuben Shaw

## Abstract

Mitophagy is a selective autophagic process that eliminates damaged mitochondria via lysosomal degradation, playing a crucial role in maintaining cellular metabolic balance. Mitophagy can occur through two pathways: ubiquitin-dependent and ubiquitin-independent. Recently, we and others have shown that, upon mitochondrial stress, AMP-activated protein kinase (AMPK) contributes to Parkin-mediated, ubiquitin-dependent mitophagy. The ubiquitin-independent pathway involves multiple outer mitochondrial membrane (OMM) “mitophagy receptors” that contain LC3-interacting region (LIR) motifs, including BNIP3, NIX/ BNIP3L, FUNDC1, and BCL2L13. LIR motifs bind Atg8/LC3 family proteins, facilitating the recruitment of the autophagosome membrane to target damaged mitochondria for degradation. The kinase Unc-51 Like autophagy activating kinase 1 (ULK1) phosphorylates the serine preceding the LIR motif in BNIP3, NIX, and FUNDC1, enhancing their binding to LC3 and promoting mitophagy. However, while BCL2L13 has been identified as a ULK1 binding partner, its regulation by phosphorylation remains unclear. We utilized mass spectrometry (MS) to map phosphorylation sites in BCL2L13 following mitochondrial stress and developed phospho-specific antibodies against two sites, Ser261 and Ser275, which were induced after exposure to the mitochondrial uncoupler, CCCP. Endogenous BCL2L13 Ser261 and Ser275 were both phosphorylated in an AMPK-dependent manner in cells and tissues. As neither site matches the established AMPK substrate consensus motif, we sought to identify which kinases directly mediate their phosphorylation downstream of AMPK. Surprisingly, genetic studies revealed that ULK1 is not regulating either site, but instead, TBK1 is controlling Ser275. This work reveals that BCL2L13 is unique amongst mitophagy receptors in being activated by mitochondrial stress and innate immune stimuli in an AMPK- and TBK1-dependent manner.

## Introduction

Mitochondria are dynamic organelles that are continuously recycled to maintain mitochondrial integrity and support critical cellular functions, including energy metabolism, stress signaling, innate immunity and even cell death. Maintaining a healthy network of mitochondria is essential to prevent pathology in humans and the accumulation of damaged mitochondria is implicated in various diseases and aging^1–4^. To prevent such buildup, cells have developed a highly conserved mechanism for mitochondrial quality control known as mitophagy. Mitophagy is a selective form of autophagy that targets dysfunctional mitochondria for degradation^5–8^. This process begins with the formation of autophagosomes, double-membrane organelles that sequester damaged mitochondria for subsequent lysosomal-mediated degradation^44,6,9^. Mitophagy is crucial for restoring energy balance, reducing mitochondrial reactive oxygen species (ROS) production, and preventing apoptosis^10,11^. Efficient removal of defective mitochondria preserves mitochondrial network integrity and safeguards against cellular damage and dysfunction.

Mitophagy can occur through ubiquitin-dependent and ubiquitin-independent pathways. The PINK1/Parkin pathway regulates ubiquitin-dependent mitophagy. When mitochondria undergo depolarization, PINK1 stabilizes on the OMM, dimerizing and triggering the recruitment of Parkin. PINK1 then phosphorylates Parkin, which subsequently ubiquitinates the mitochondria, marking them for degradation^12–15^. Recently, our lab has shown that AMPK also plays an early role in PINK1/Parkin-mediated mitophagy, triggering phosphorylation of Ser108 of Parkin in the cytoplasm within 5 minutes of CCCP treatment, which precedes subsequent phosphorylation of Parkin on Ser65 by PINK1 at the mitochondria at 30 to 60 minutes^16^. The regulation of Parkin Ser108 by AMPK was also corroborated in a recent study that further clarified the roles of multiple kinases in different mitophagy contexts^17^.

Ubiquitin-independent mitophagy involves several OMM proteins, such as NIX, BNIP3, FUNDC1, and BCL2L13. These proteins contain LC3-interacting region (LIR) motifs, which directly bind to Atg8 proteins, facilitating the recruitment of mitochondria into autophagosomes for mitophagic processing^18–23^. Remarkably, of the ∼50 mammalian proteins containing a LIR motif, less than ten of them contain a serine preceding the first residue of the LIR motif, yet all four aforementioned mitophagy receptors bear a serine in this position (FUNDC1, BNIP3, NIX, and BCL2L13)^24,25^. Multiple biochemical, cellular, and structural studies have found that phosphorylation of the serine flanking the LIR motif enhances the binding of mitophagy receptor proteins to the Atg8 proteins, promoting the selective sequestration of mitochondria into autophagosomes for degradation^24,26–28^.

Initial studies of p62/SQSTM1 ^29–31^ and other Sequestrosome-like Receptors (“SLR”) OPTN and NDP52^14,32,33^ revealed their phosphorylation on the serine preceding their LIR motif by the kinase TBK1, which has well-established roles in multiple forms of autophagy, including mitophagy ^34–37^. Recent studies identified ULK1 as the kinase responsible for phosphorylating the Serine preceding the LIR in three of the mitophagy receptors: FUNDC1, BNIP3, and NIX^27,38^. However, while BCL2L13 has been characterized as a binding partner of ULK1, the specific kinase(s) involved in its regulation remain unclear. BCL2L13, also known as BCL Rambo^39^, is a unique member of the BCL2 family of proteins.

The BCL2 family is primarily recognized for its role in regulating apoptosis, with members classified into pro-survival (e.g., BCL2, BCL-XL) and pro-apoptotic (e.g., BAX, BAK) proteins^40,41^. Like some BCL2 family members, BCL2L13 contains four BH domains (BH1, BH2, BH3, and BH4), which are typically associated with apoptosis and mitochondrial fragmentation. However, unlike canonical BCL2 proteins, BCL2L13 functions as a mitophagy receptor. In contrast, NIX and BNIP3 are considered atypical BCL2 family proteins, as they contain only a single BH3 domain and primarily facilitate mitophagy, particularly under hypoxic conditions. While BCL2L13 was initially thought to function primarily in apoptosis, emerging evidence suggests that it plays a role in mitophagy^41–43^. As a ubiquitin-independent mitophagy receptor, BCL2L13 is homologous to budding yeast ATG32 and, notably, can functionally replace it^44^. Like ATG32, BCL2L13 contains a canonical LIR motif and a C-terminal transmembrane domain that anchors it to the OMM. It also interacts with ULK1^45^, paralleling ATG32, which binds the ULK1 yeast homolog ATG1^44^. Despite these similarities, it remains unclear whether ULK1 phosphorylates BCL2L13 in mammalian cells or which specific cellular stresses activate BCL2L13-dependent mitophagy. In contrast, NIX, BNIP3, and FUNDC1 are all established to be phosphorylated by ULK1, and all three were reported to be activated under hypoxic conditions^38,46–48^.

We aimed to investigate whether BCL2L13 is regulated by phosphorylation and which kinases are involved. To our surprise, we discovered that BCL2L13 is regulated by a novel dual mechanism involving AMPK and TBK1, rather than ULK1. Specifically, mitochondrial damage or direct AMPK activators promote phosphorylation of endogenous BCL2L13 at serine 261 and serine 275 in mouse embryonic fibroblasts (MEFs) and mouse liver. Additionally, BCL2L13 plays a role in CCCP-induced mitophagy in cells lacking the other mitophagy receptors, highlighting the compensatory roles these mitophagy receptors play in mitophagy and identifying areas for future research.

## Results

### Mitochondrial OXPHOS inhibition or direct AMPK activators induce phosphorylation of BCL2L13

Phosphorylation of serines preceding LIR sequences was first reported in the few SLR receptors that contain a serine directly preceding their LIR motifs, including P62/SQSTM1, NDP52, and OPTN. Phosphorylation of serines adjacent to LIR sequences has been shown to increase LC3-binding, providing a stimulus-specific mechanism for induced LC3-interactions^3,33,35,49^. A common characteristic of the ubiquitin-independent mitophagy receptors is that most of the well-characterized receptors (BNIP3, NIX/BNIP3L, FUNCDC1, BCL2L13) share a serine in front of their defined LIR motifs, regardless of where the LIR motif was located within the broader protein structure (**Fig. 1A**). Indeed, over the last 5 years, FUNDC1, BNIP3, and NIX have all been reported to be phosphorylated on these LIR-preceding serines by the autophagy kinase ULK1, and this event has been suggested to activate these mitophagy receptors following hypoxia^17,50^. This led us to examine the regulation of BCL2L13 by phosphorylation after mitophagy-inducing stimuli, especially since it has already been suggested to directly bind ULK1^45^. To identify stimuli regulating BCL2L13 phosphorylation, we treated U2OS cells stably expressing flag-tagged BCL2L13 with the mitochondrial oxidative phosphorylation (OXPHOS) uncoupler CCCP, the mitochondrial Complex I inhibitor rotenone, the direct AMPK activator compound 991, the hypoxia mimetic DFP, or the mTORC1 catalytic inhibitor INK128. Following a 1-hour treatment or each of these, we immunoprecipitated BCL2L13 using anti-flag beads and performed mass spectrometry phosphoproteomics to map phosphorylation sites on BCL2L13 (**Fig. S1A,B**). We found that treatment with CCCP and compound 991 induced more phosphorylation sites on BCL2L13 compared to DFP and INK128 (**Fig. S1C**). Of the novel identified phosphorylation sites, serine 259 and serine 261 aligned with the consensus motifs of ULK1^51^ and TBK1^52^, but not with those of AMPK^53^. A deeper analysis of the basal and CCCP-induced phosphorylation sites in BCL2L13 revealed their enrichment in a 25 amino-acid stretch preceding the LIR motif, which interestingly contains two nearly identical serine-rich repeat regions (**Fig. 1B**).

**Figure 1.**
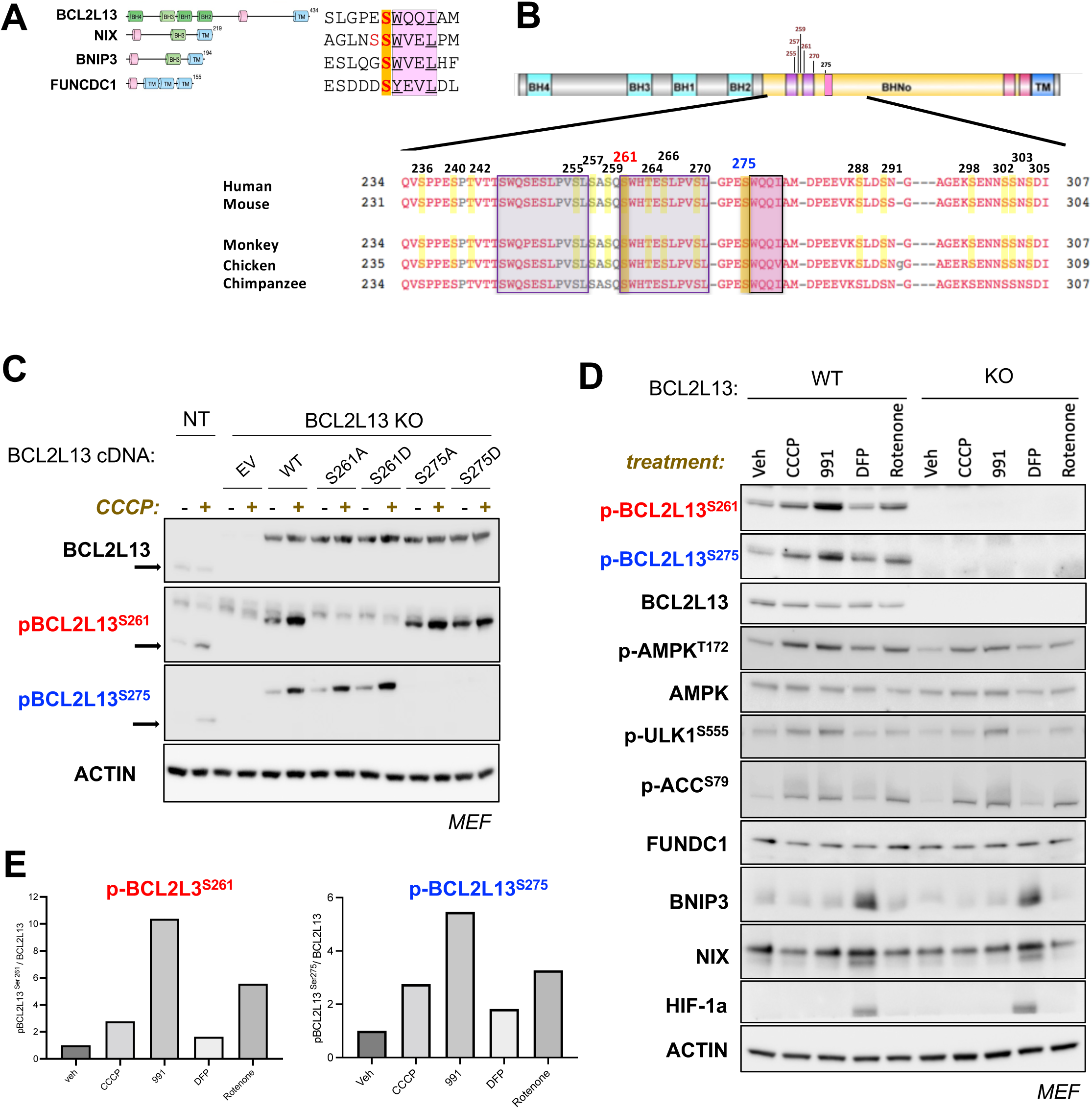
Identification of Ser261 and Ser275 as BCL2L13 phosphorylation sites induced by CCCP. **(A)** Schematic representation of the domain structures of mitophagy receptor proteins, highlighting the LIR motifs (pink color) and the C-terminal transmembrane (TM) domain (blue color). **(B)** Clustal Omega alignment of BCL2L13 orthologs across species showing conservation of reported phosphorylation sites. Note that S261 lies at the start of the second of two repeat elements (purple hue) which precede S275 and the LIR motif (pink hue). Reported phosphorylation sites in mass spectrometry datasets are highlighted in yellow. **(C)** Immunoblots of WT (NT) or CRISPR/Cas9 RNP-BCL2L13 knockout (KO) MEF cell lines bearing stable lentiviral expression of indicated BCL2L13 cDNAs, treated with 20 µM CCCP for 1 hour. These blots demonstrate that the phospho-specific antibodies detect BCL2L13 phosphorylation specifically at serine 261 (S261) and serine 275 (S275) in MEF cells, with phosphorylation levels increasing following CCCP treatment. Arrows indicate endogenous mouse Bcl2l13 (predicted to be 7 kDA smaller versus human). **(D)** Phosphorylation of endogenous Bcl2L13 on Ser261 and Ser275 compared to the regulation of other mitophagy receptor protein levels. Immunoblots with indicated antibodies on WT or BCL2L13 KO MEF cells, treated with DMSO vehicle (veh) or 10 µM 991 or 20 µM CCCP or 1 µM rotenone or 1mM DFP. **(E)** Quantification of BCL2L13 Ser^261^ (left) and BCL2L13 Ser^275^ (right) from the immunoblot shown in panel D. Densitometric analysis was performed to assess protein expression levels. Phospho-protein levels were quantified and normalized to total protein levels.

To further study BCL2L13 regulation by phosphorylation, we developed two distinct phospho-specific antibodies against P-Ser261 and P-Ser275. In order to study the specificity of our phospho-specific antibodies, we created cell lines using ribonucleoprotein (RNP) complex oligonucleotide sgRNA guides with CRISPR/Cas9 to genetically disrupt endogenous BCL2L13. This was confirmed by immunoblotting with an antibody we validated as capable of recognizing endogenous levels of BCL2L13 in these cells (**Fig. S3F**). We then stably introduced lentiviral vectors expressing cDNAs encoding wild-type (“WT”), non-phosphorylatable (“S261A”, “S275A”) or phospho-mimetic (“S261D”, “S275D”) forms of BCL2L13. We then subjected this isogenic cell panel to CCCP or DMSO (vehicle control) treatment and immunoblotted with antibodies we characterized against total BCL2L13, P-Ser261 BCL2L13, or P-Ser275 BCL2L13. Phosphorylation of both Ser261 and Ser275 increased following CCCP in the endogenous and tagged WT BCL2L13, but not in the tagged serine-mutant BCL2L13 proteins (**Fig. 1C**).

Next, we tested a broader panel of mitophagy-inducing stimuli and examined endogenous BCL2L13 phosphorylation using our validated phospho-specific antibodies. Phosphorylation of serine 261 and serine 275 increased after CCCP treatment, along with activation of AMPK targets, including phospho-ACC at serine 79 and phospho-ULK1 at serine 555^54^ (**Fig. 1D, E**). We also tested compound 991, a small molecule that directly binds to and allosterically activates AMPK, independent of changes in mitochondria or cellular energy levels^55^. Treatment with 991 resulted in a significant increase in the phosphorylation of BCL2L13 at serine 261 and serine 275 (**Fig. 1D, E**). Lastly, treatment with the mitochondrial complex I inhibitor rotenone induced phosphorylation of BCL2L13 at serine 261 and serine 275 (**Fig. 1D, E**).

In contrast, the hypoxic mimetic iron chelator deferiprone (DFP) did not induce phosphorylation of BCL2L13 at either serine 261 or serine 275, consistent with our earlier phospho-proteomic analysis (**Fig. S1A-C**). As expected, DFP increased levels of hypoxia-inducible factor (HIF-1α), BNIP3, and NIX in these cells (**Fig. 1D**), consistent with the binding of HIF-1α to the promoters of BNIP3 and NIX and increasing their transcription^50^. None of these stimuli resulted in consistent changes in FUNDC1 or BCL2L13.

### BCL2L13 serine 261 and serine 275 phosphorylation occur in *vivo* upon AMPK activation

Having established that BCL2L13 is phosphorylated in MEFs at serine 261 and serine 275 by direct AMPK activators (991) and mitochondrial OXPHOS inhibitors that activate AMPK, we next directly tested whether phosphorylation of BCL2L13 at S261 or S275 is genetically dependent on AMPK in mouse liver, a central metabolic organ whose homeostasis is highly regulated by AMPK^56^. We assessed protein levels by immunoblotting liver lysates from control and AMPKα1/α2 liver knockout (AMPK DKO) mice treated with either vehicle or the AMPK activator MK8722 for 2 hours. In wild-type (WT) mouse livers, BCL2L13 phosphorylation at serine 261 and serine 275 increased after MK8722 treatment. These phosphorylation events were absent in AMPK DKO livers, consistent with changes observed in well-known AMPK substrates, such as phospho-ACC and phospho-ULK1 (**Figure 2A, S2A**). Previous studies have hinted that the BCL2L13 Ser275 preceding the LIR is regulated by ULK1, as ULK1 has been established to bind BCL2L13^45^, and ULK1 has been reported to phosphorylate the serine preceding the LIR in all of the other UB-independent mitophagy receptors. We next directly tested whether phosphorylation of BCL2L13 at S261 and S275 is genetically dependent on ULK1 and its close homolog ULK2, using liver lysates from control and liver-specific ULK1/2 double knockout (ULK DKO) mice treated with either vehicle or MK8722. BCL2L13 phosphorylation at S261 and S275 increased following AMPK activation, even in the absence of ULK1/2 (**Fig. 2B, S2A**).

**Figure 2.**
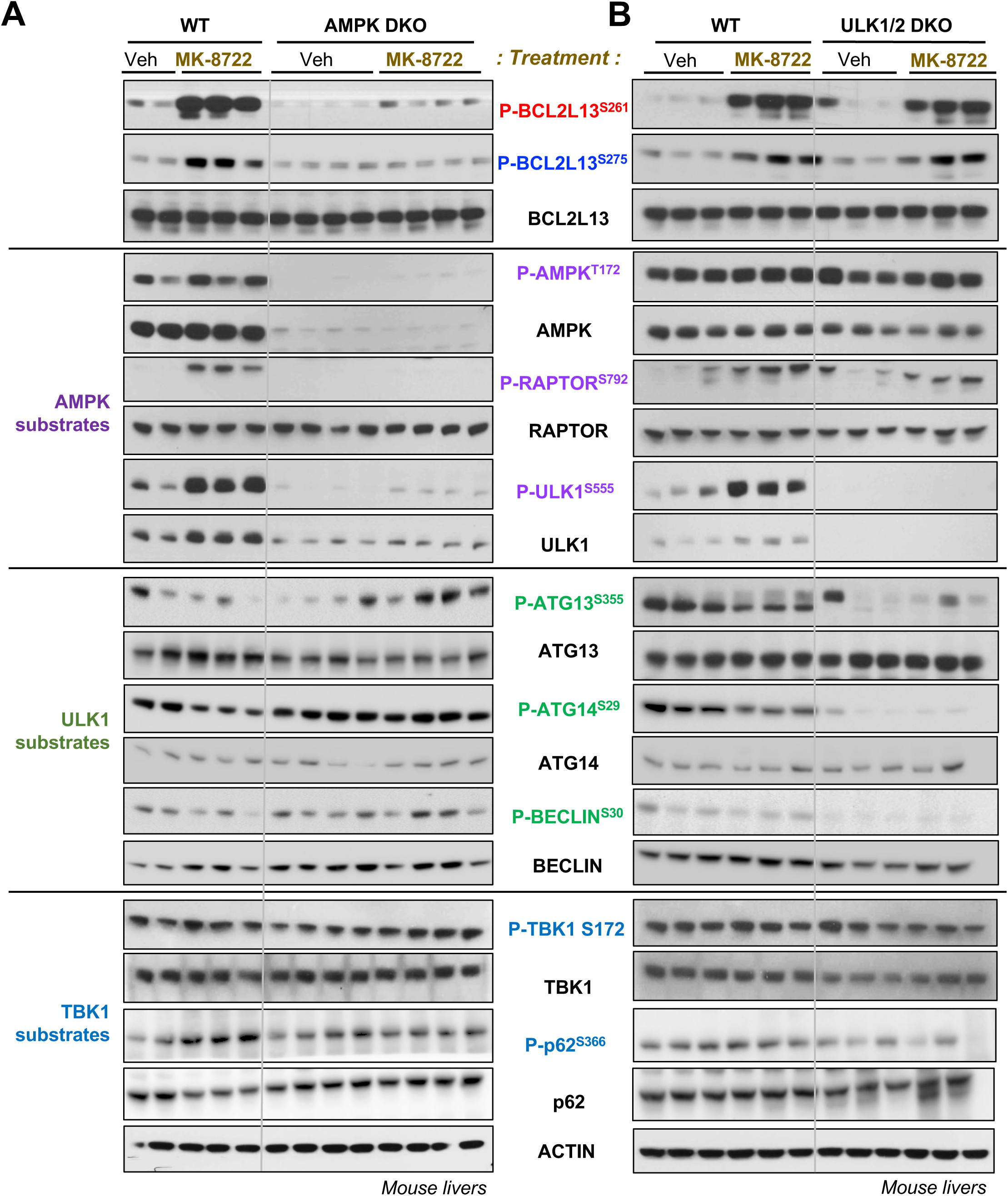
Endogenous BCL2L13 phospho-Ser^261^ and Ser^275^ are AMPK-dependent but ULK1/2-independent in mouse liver. (**A**) Immunoblots of liver lysates from control or inducible knockout of AMPKa1/a2 (AMPK DKO) mice gavaged with vehicle (veh) or 30mg/kg AMPK activator MK-8722 for 2 hours. **(B)** As in panel A, except from control or knockout of ULK1 and ULK2 (ULK1/2DKO). Immunoblots in panel A and B probed with antibodies indicated in the center. In wild-type (WT) mice, MK8722 treatment resulted in increased phosphorylation of AMPK substrates, including pRaptor^S792^, and pULK1^S555^, all of which are completely abolished in the AMPK DKO lysates. In contrast, MK8722 treatment in wild-type livers results in a modest decrease in the phosphorylation of ULK1 substrates, such as pATG13^S355^, pATG14^S29^, and pBeclin^S30^. This reduction in ULK1 substrate phosphorylation after MK8722 appears diminished in the AMPK DKO livers, suggesting that AMPK may be inhibiting ULK1 kinase activity towards these substrates. The phosphorylation of ATG13, ATG14, and Beclin on the indicated sites is fully ablated in the ULK1/2 DKO, reinforcing the specificity of the antibodies and verifying that these are indeed ULK1/2 phosphorylation sites in vivo. Quantification of all these sites compared to their total protein levels is seen in Supplemental Fig. 2.

Consistent with Ser261 and Ser275 not being ULK1/2 sites, the phosphorylation of three previously reported ULK1 substrates (Atg13 Ser355, Atg14 Ser29, Beclin Ser30) was distinctly reduced following MK8722 treatment in WT livers (**Fig 2A, S2A**), and this effect was attenuated in the AMPK KO livers, consistent with recent reports that AMPK phosphorylation of ULK1 leads to an inhibition of ULK1 kinase activity, while stabilizing ULK1 levels via 14-3-3 binding^17^. The phosphorylation of these three reported ULK1 sites (Atg13 Ser355, Atg14 Ser29, Beclin Ser30) was abolished in the ULK1/2 DKO livers, confirming both the specificity of the antibodies and that these residues are ULK1/2 substrates in vivo. This also indicates that no other kinase can compensate and target these same sites when ULK1/2 is genetically ablated. TBK1 is another kinase that is well-established to be involved in mitochondrial stress-induced mitophagy^57–59^, and TBK1 has also been reported to be stimulated by AMPK phosphorylation in some contexts^60^. Given that TBK1 has a similar kinase motif specificity to ULK1, and TBK1 has been shown to phosphorylate -1 LIR Serines in p62, TAXBP1, and OPTN^29,33,61,62^, we examined readouts of TBK1 activity in our WT, AMPK, and ULK1 KO livers. MK8722 treatment modestly increased P-S172 TBK1 and phosphorylation of the reported TBK1 site Ser366 in p62/SQSTM1 in WT livers but not AMPK KO livers (**Fig. 2A, S2A**). An antibody was not available to the previously reported AMPK phosphorylation site in TBK1, Ser511^60^. It is worth noting that TBK1 activity appeared increased basally in the ULK1/2 DKO livers, consistent with previous reports of TBK1 and TAXBP1 upregulation in genetic knockouts of other ULK-complex components ^63^.

Our findings in Fig. 2 are partially consistent with a recent report that used kinase siRNA libraries to identify kinases regulating BCL2L13 S275, where AMPKα2 was found^64^. However, neither serine 261 nor serine 275 aligns with the well-established AMPK consensus motif^53^, suggesting an intermediate kinase likely mediates their phosphorylation. Furthermore, a whole kinome library of optimal substrate specificity was recently reported for all human kinases^57^, which we utilized to predict the top 20 likely kinases for BCL2L13 Ser261 or Ser275 (**Fig. S2B)**. AMPK was not predicted in the top 20 likely kinases for either site (**Fig. S2B**), consistent with its lack of the known amino acids surrounding the phosphorylation site that confer AMPK specificity. Notably, TBK1 was the top predicted kinase to phosphorylate Ser275 based on its amino acid sequence (**Fig. S2B**).

### BCL2L13 S275 is phosphorylated by TBK1 in MEFs

Given the possibility that TBK1 may be an intermediate kinase between AMPK and BCL2L13, we next directly tested whether TBK1 was required for BCL2L13 S261 and S275 phosphorylation. We used CRISPR-Cas9 RNP to delete TBK1 in MEFs and compared BCL2L13 S261 and S275 phosphorylation following treatment with the classic TBK1 activating stimulus Lipopolysaccharide (LPS). BCL2L13 S275 phosphorylation increased following LPS treatment in WT, but not TBK1 knockout (KO) MEFs (**Fig. 3A**). In contrast, BCL2L13 phosphorylation at S261 remained unaffected in these cells. To further investigate the observed dual regulation of BCL2L13, we examined its phosphorylation at S261 and S275 in WT, TBK1 KO, and AMPK DKO MEFs treated with LPS, CCCP, or MK8722. As seen previously, phosphorylation of BCL2L13 at S261 increased following treatment with CCCP and MK8722, and this increase was attenuated in AMPK DKO MEFs (**Fig. 3B, 3C**). Treatment with LPS did not affect BCL2L13 S261 phosphorylation. In contrast, phosphorylation of BCL2L13 at S275 was elevated after LPS, CCCP, and MK8722 treatments in WT cells, but this increase was abolished in TBK1 KO cells (**Fig. 3B, 3C**). Additionally, while MK8722-induced BCL2L13 S275 phosphorylation was lost in AMPK DKO cells, the increase in BCL2L13 S275 by LPS and CCCP was not reduced in AMPK DKO cells. This suggests that, while direct AMPK activation can stimulate TBK1, other well-established stimuli, LPS and CCCP, can lead to TBK1 activation via TRIF-TRAF3^65^ and OPTN-NDP52^33,57^, respectively, without a requirement for AMPK phosphorylation of TBK1. Regardless of which stimulus activates TBK1 in these cells, it was always paralleled by increases in BCL2L13 Ser275 phosphorylation.

**Figure 3.**
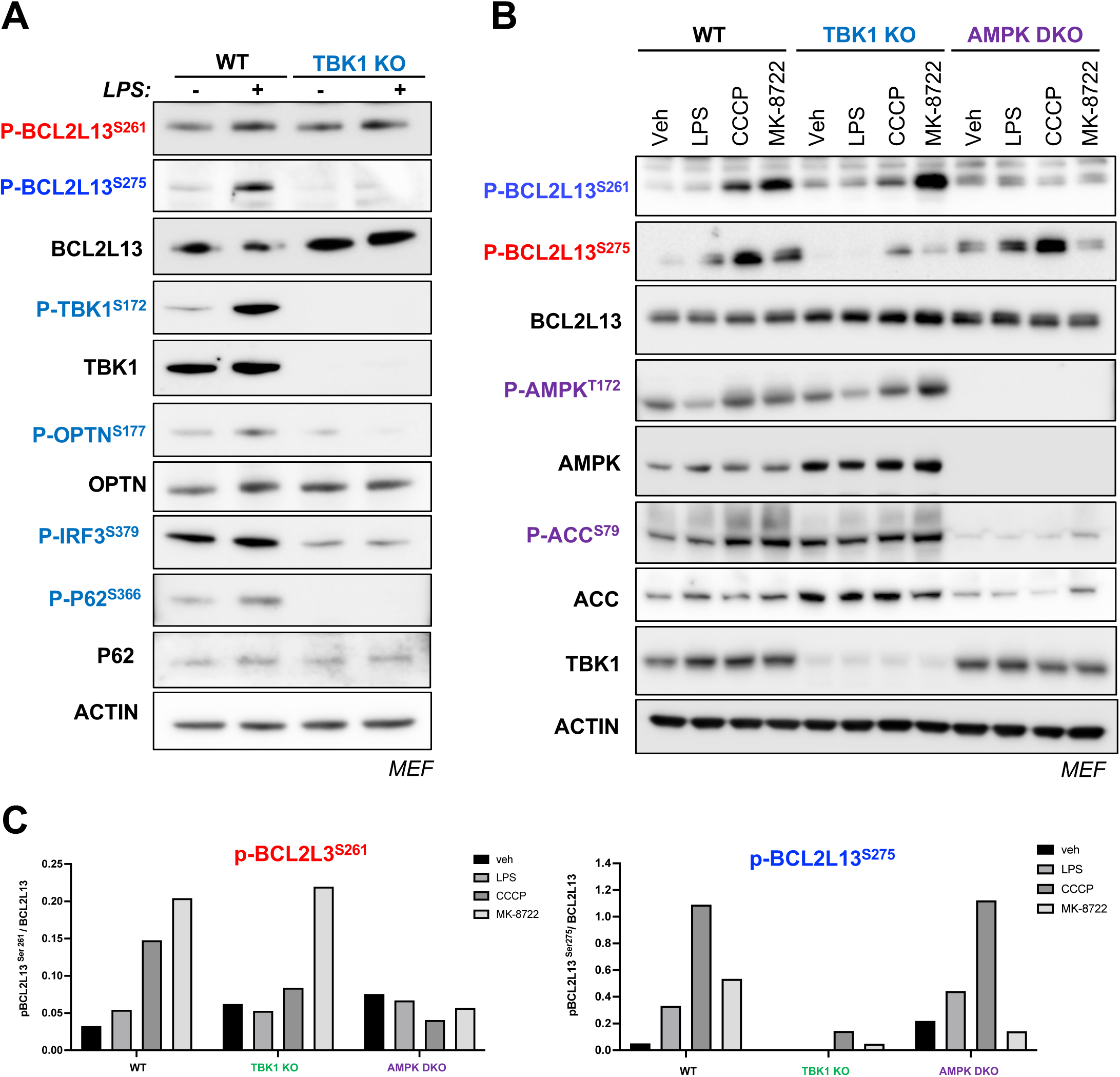
BCL2L13 Ser^275^ is regulated by TBK1. **(A)** Immunoblots of MEF cells from wild-type (WT) or CRISPR-Cas9 RNP-mediated TBK1 knockout (KO) lines, treated with vehicle or TBK1 activator LPS for 1 hour. **(B)** Immunoblot analysis of wild-type (WT) MEFs, TBK1 knockout (KO) MEFs, and AMPK knockout (AMPKDKO) MEFs treated with vehicle (Veh), LPS, CCCP, or MK8722. **(C)** Quantification of BCL2L13 Ser^261^ (left) and BCL2L13 Ser^275^ (right) from the immunoblot shown in B. Densitometric analysis was performed to assess protein expression levels. Phospho-protein levels were quantified and normalized to total protein levels and loading control, Actin.

### Genetic deletion of BCL2L13 does not impact CCCP- or DFP-induced mitophagy in MEFs

Previous RNAi studies in HEK293A cells suggested that BCL2L13 depletion dampens CCCP-induced mitophagy, though this was assessed indirectly by counting LC3B and mitochondrial ATP synthase double-positive puncta^66^. In addition, another RNAi study in ear mesenchymal stem cells from MitoQC mice suggested that knockdown of BCL2L13 decreased DFP-induced mitophagy, as measured by immunofluorescence (IF) quantifying mCherry puncta^67^. Based on these prior studies, we chose to explore the functional role of BCL2L13 in mitophagy using HEK293A cells and MEF cells, which are more similar to murine ear mesenchymal cells than other readily available cell lines. We first generated MEF and HEK293A cells that stably express the mitophagy reporter, Mito-QC (**Fig. S3A**). This reporter is a GFP-mCherry fusion protein targeted to the outer mitochondrial membrane (OMM) via a FIS1-targeting sequence. Under physiological conditions, cells express equal ratios of GFP and mCherry signals. However, when mitochondria are targeted to lysosomes, the acidic environment within the lysosome quenches the pH-sensitive GFP signal increasing the ratio of mCherry to GFP^50^. Therefore, an increase in the mCherry signal serves as a readout of active mitophagy, which can be quantified by flow cytometry or fluorescence microscopy (**Fig. S3B**). Next, we established the optimal time points to observe maximal CCCP- and DFP-induced mitophagy in our WT MEF MitoQC (“MEF-QC”) and HEK293A MitoQC (“HEK293A-QC”) cells. In MEFs, mitophagy peaked between 2-6 h after CCCP treatment and reduced to baseline after 18-24 h (**Fig. S3C**). In contrast, DFP-induced mitophagy peaked at 18-24h in MEFs (**Fig. S3C**), consistent with the known transcriptional induction of BNIP3 and NIX by HIF-1α. We chose a 4h CCCP and a 24h DFP timepoint for subsequent analyses in MEFs. In contrast, in HEK293A cells, the MitoQC signal peaked at 6h in response to both CCCP and DFP (**Fig. S3D**), so we used that timepoint for subsequent analysis in these cells. Immunoblot analysis of these timepoints in MEFs and HEK293A cells (**Fig. S3E**) revealed HIF-1α increases after DFP (but not CCCP) at 2h, 4h, and 6h in both cell types, with changes in BNIP3 or NIX after DFP more apparent at 18h (note the BNIP3 antibody used did not react with mouse BNIP3). Modest increases were observed in BCL2L13 levels in both cell lines at 2h and 4h after CCCP, whereas clear decreases in FUNDC1 levels after CCCP were observed in both cell lines (**Fig. S3E**).

Having established the optimal timepoints to examine DFP- and CCCP-induced mitophagy as readout by the MitoQC reporter in both these cell types, we proceeded to make CRISPR/Cas9 RNP-mediated knockouts of BCL2L13 in the MEF-QC and HEK293A-QC cells (**Fig. S3F**). In both WT MEF-QC and WT HEK293A-QC, CCCP and DFP induced significant mitophagy, as measured by the increased mCherry/GFP ratio. Notably, no difference in mitophagy was observed between WT and BCL2L13 KO cells following either CCCP or DFP treatment under the conditions we used (**Fig. S3G-S3H)**. These findings suggest that genetic deletion of BCL2L13 alone does not significantly impact CCCP or DFP-induced mitophagy in MEF or HEK293A cells.

We next explored whether we observed any sign of compensation for loss of BCL2L13 as evidenced by upregulation of other mitophagy receptors in these cells. As seen in **Fig. S3I** and **Fig. S3J**, we saw modest increases in NIX in the BCL2L13 knockout MEF-QC and HEK-293QC cells. In contrast to HEK293A, MEF cells displayed greater differential regulation of BCL2L13 levels, with increases after CCCP and decreases after DFP (compare **Fig. S3I** to **Fig. S3J**). Based on these observations, we decided to focus our analysis to MEF cells for further investigations on the role of BCL2L13 in mitophagy.

### CRISPR deletion of four Ubiquitin-independent mitophagy receptors reduces CCCP-induced mitophagy in MEFs

Given that BCL2L13 deletion alone does not affect CCCP- or DFP-induced mitophagy, and that BCL2L13 phosphorylation is increased after CCCP, but not DFP treatment, we further focused on CCCP-induced mitophagy. Specifically, we aimed to investigate whether other mitophagy receptors compensate for the loss of BCL2L13 during CCCP-induced mitophagy. Using CRISPR-Cas9, we generated various knockout (KO) cell lines in the MEF-QC cells. These included single knockouts of BCL2L13, NIX, BNIP3, or FUNDC1, as well as a triple KO (NIX, BNIP3, and FUNDC1) and a quadruple KO (all four receptors: BCL2L13, NIX, BNIP3, and FUNDC1) (**Fig. 4A**). The cells were then treated with CCCP and mitophagy was analyzed using flow cytometry. None of the individual mitophagy receptor KO cells had an impact on CCCP-induced mitophagy under the conditions we used. In fact, all of them showed modest increases in mitophagy, an effect lost when BNIP3, NIX, and FUNDC1 knockouts were combined (“Triple KO”) – see black bars in **Fig. 4B**. However, the Triple KO cells did not exhibit a statistically significant reduction in their ability to undergo mitophagy compared to the control WT MEF-QC cells. In contrast, cells lacking all four receptors (QUAD KO) exhibited a significant reduction in CCCP-induced mitophagy, compared to the WT MEF-QC cells, and when comparing the Triple KO to the Quad KO (**Fig. 4B**). These results indicate that loss of BCL2L13 is specifically required for mitophagy in cells lacking BNIP3, NIX, and FUNCDC1.

**Figure 4.**
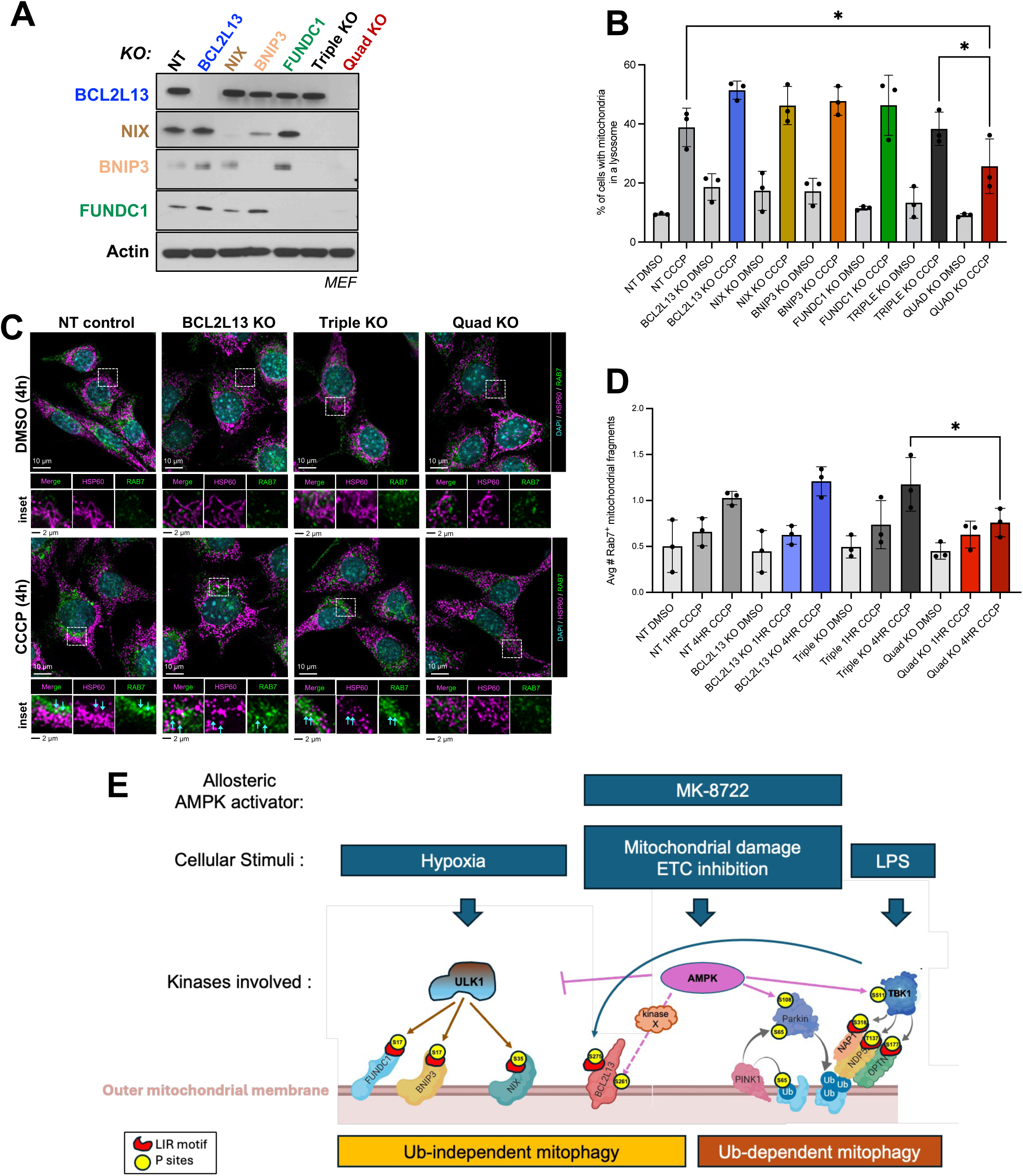
Redundant roles for Ubiquitin-independent mitophagy receptors in CCCP-induced mitophagy. **(A)** Immunoblot analysis of CRISPR/Cas9 knockouts of individual mitophagy receptors, and triple and quadruple deletions in MEFs. **(B)** MitoQC functional mitophagy assay after CCCP in MEFs from panel A, as analyzed by flow cytometry. Quantitative analysis was performed on the ratiometric flow cytometry data (mCherry/GFP). Data are presented as mean ± SEM (n = 3). Statistical significance was determined using one-way ANOVA. Asterisks indicate significant differences compared to the control group (\**p* < 0.05). **(C)** Representative confocal micrographs of Control (NT), BCL2L13 KO, Triple KO, and Quad KO MEFs treated with 20 µM CCCP for the indicated timepoints and immunostained for HSP60 and Rab7. Zoomed-in regions highlight co-localization of HSP60 and Rab7 (indicated by blue arrows). **(D)** Mitolysosome formation was quantified as RAB7+ mitochondrial fragments using CellProfiler. Data points represent experimental averages of more than 200 cells per condition (n = 3). Statistical significance was determined using one-way ANOVA. Asterisks indicate significant differences compared to the control group (\**p* < 0.05). **(E)** Model for the regulation of BCL2L13. After mitochondrial damage, AMPK activates Parkin and BCL2L13, while simultaneously suppressing NIX and BNIP3 activation by ULK1. In contrast, under hypoxic conditions, ULK1-dependent phosphorylation and activation of BNIP3, NIX, and FUNDC1 are critical events, and neither Parkin nor BCL2L13 appears to be involved. Thus, Bcl2L13 is currently the only ubiquitin-independent mitophagy receptor known to be acutely activated by mitochondrial damage.

To independently examine the role of BCL2L13 in mitophagy, we also performed immunofluorescence (IF) analysis to quantify mitolysosome formation in the same receptor knockout backgrounds. We defined mitolysomes as cells bearing quantitative co-localization of the mitochondrial marker HSP60 and the late endosomal marker RAB7. In order to perform this analysis, we needed cells lacking the MitoQC reporter, so we remade BCL2L13 KO alone, Triple KO, and Quadruple KO in WT MEFs without the reporter. Consistent with the flow cytometry MitoQC results, quantitative high-throughput indirect immunofluorescence of mitolysosome formation in the WT, BCL2L13 KO, Triple KO and Quad KO MEFs revealed a marked reduction in mitolysomes in the Quad KO compared to the triple KO. These results reinforce the critical role of BCL2L13 in CCCP-induced mitophagy in the MEFs lacking BNIP3, NIX, and FUNDC1 (**Fig. 4C and 4D**). As observed in the flow cytometry MitoQC cell analyses, no difference in mitolysome formation after CCCP was seen in the BCL2L13 KO alone or Triple KO MEF-cells when compared to the WT MEF cells (**Fig. 4D**). Only the Quad KO cells exhibited a significant loss of HSP60/RAB7 colocalization after CCCP, indicating less mitophagy.

Finally, we sought to use the Quad KO MEFs to investigate the functional requirement of BCL2L13 phosphorylation at S261 and S275 in CCCP-dependent mitophagy. We therefore stably reintroduced a WT or Serine-to-Alanine (SA) mutant flag-tagged BCL2L13 cDNAs into Quad KO MEF-QC cells or Quad KO MEF cells to perform flow cytometry MitoQC analysis or HSP60/RAB7 mitolysosome quantitative colocalization analysis as described above. Despite equivalent levels of WT, S261A, or S275A mutant expression (**Fig. S4A**), we did not observe restoration of functional mitophagy in the Quadruple KO cells with WT BCL2L13 added back (**Fig. S4B, C**). Comparison of S261A or S275A mutants to WT also revealed no significant differences in mitophagy rates across assays or cell lines (**Fig. S4B, C**).

## Discussion

This study has revealed a number of novel findings regarding the role of BCL2L13 in CCCP-dependent mitophagy. We uncovered BCL2L13 Ser261 and Ser275 as novel phosphorylation sites that are dependent on AMPK in cells in culture and mouse liver in vivo. Ser261 is a previously undescribed phosphorylation site that is located roughly fifteen amino acids upstream of the LIR motif at the start of a unique repeated sequence of unknown function in BCL2L13 (**Fig. 1B**). Across cells and conditions, Ser261 was tightly associated with AMPK activity, mirroring well-established AMPK phosphorylation sites like Raptor Ser792 and ULK1 Ser555. The Ser261 phosphorylation increase was ablated in the absence of AMPK, but remained unaffected by the absence of TBK1. In contrast, Ser275, which is the serine directly preceding the LIR motif, is regulated by TBK1 following mitochondrial uncoupling by CCCP or TBK1 activation by LPS. Phosphorylation at serine 275 was reduced in the absence of TBK1 regardless of stimulus. These findings indicate that BCL2L13 is phosphorylated on distinct sites in response to different upstream stimuli. Notably, neither of these sites is dependent on the one reported binding partner of BCL2L13, ULK1^45^, despite ULK1 being a predicted candidate kinase for Ser261 based on the amino acid sequence flanking Ser261 (**Fig. S2B**). This suggests the existence of an unknown kinase whose activity is stimulated by AMPK and which may mediate direct phosphorylation of Ser261. Kinases predicted by the optimal kinome library approach^68^ in **Fig. S2B** represent good candidates for further exploration.

Recently, Ser275 of BCL2L13 was reported as being phosphorylated by AMPK directly and playing a critical role in mitophagy in pressure-overload in cardiac tissues in mice^64^. In that study, AMPK was identified by an RNAi screen for kinases controlling Ser275 phosphorylation, consistent with our results here. However, based on extensive analysis of AMPK substrates and TBK1 substrates, it appears most likely that AMPK cannot directly phosphorylate Ser275 in vivo, and the previously described AMPK activation of TBK1^60^ is controlling this event in cells and tissues.

Our analysis of AMPK signaling in vivo in mouse liver following the small molecule AMPK activator MK-8722 contributes to a deeper understanding of the interplay between AMPK, TBK1, and ULK1, which has become an area of greater interest in the last few years. Indeed, despite initial models based on genetics that suggested that AMPK phosphorylation of ULK1 would lead to its activation, as mTORC1 phosphorylation of ULK1 leads to its inhibition, recent studies using more specific tools and new phospho-specific antibodies against downstream ULK1 substrates have revealed a more complex relationship. Kim and colleagues first demonstrated that AMPK phosphorylation of ULK1 resulted in a suppression of ULK1 kinase activity towards multiple of its substrates (Beclin, Atg14, Atg16L1), a set of findings we now corroborate here in vivo using WT, AMPK DKO, and ULK DKO livers. Phosphorylation of ATG13 Ser355, ATG14 Ser29, and Beclin Ser30 were abolished in these endogenous proteins in the ULK1/2 DKO livers compared to the WT livers, verifying their behavior at ULK1/2 specific substrates in this tissue. These same sites are reduced in WT livers treated with the small molecule AMPK activator MK-8722, and this reduction in their phosphorylation was lost in the AMPK DKO livers. This is very clear evidence in support of the model of Kim and colleagues, consistent with a full revision of the assumed role of AMPK to activate ULK1 kinase activity. Our findings are also consistent with a more recent study from Ganley and colleagues with direct relevance to UB-independent mitophagy receptors. In their study, AMPK activation suppressed ULK1-dependent activation of BNIP3 and NIX-dependent mitophagy^17^. As ULK1 is known to directly phosphorylate the serine in front of the LIR in BNIP3 (Ser17), NIX (Ser35), and FUNCDC1 (Ser14), we surmise that AMPK activation would suppress mitophagy induced by these receptors, while simultaneously increasing phosphorylation of Ser275 in BCL2L13 in front of its LIR. Indeed, Ganley and colleagues made a model in which AMPK promotes mitophagy specifically of damaged mitochondria, while sparing mitophagy of healthy, functional mitochondria. AMPK phosphorylation of Parkin Ser108, which we previously reported ^16^ and Ganley corroborated^17^, contributes to Parkin activation and removal of damaged mitochondria. The findings here that AMPK also may activate BCL2L13 in mitophagy via Ser261 and Ser275 phosphorylation make BCL2L13 the only UB-independent mitophagy receptor known to date to play a role in CCCP-induced mitophagy. Whether the implications of these recent studies all hold up remains to be seen across many studies of additional mitophagy contexts, but the development of phospho-specific antibodies against endogenous BCL2L13 in human and mouse should facilitate this. Additional development against the aforementioned ULK1-phosphorylation sites in BNIP3, NIX, and FUNDC1 would greatly aid the field to clarify their individual and combined roles in different tissues and pathological conditions.

In contrast to the elegant work from Murakawa and colleagues using a Ser275Ala knock-in mouse in which specific cardiac defects in mitophagy were observed^64^, we could not observe strong effects on CCCP-induced mitophagy from altered BCL2L13 in MEFs or HEK293A cells in culture, unless BNIP3, NIX, and FUNDC1 were also co-deleted. Indeed, all of these mitophagy receptors were appreciably expressed in MEFs and HEK293A cells, and CCCP treatment may also represent a large non-physiological induction of mitochondrial damage, which cannot parse out the requirements for endogenous mitophagy receptor functions and post-translational regulations. Our analysis with single knockouts of BCL2L13, NIX, BNIP3, or FUNCDC1 revealed that none of them alone are required for CCCP-induced mitophagy in MEFs under our conditions to acutely maximize mitophagy; indeed, even triple KO of BNIP3, NIX, and FUNCDC1 had minimal impact on MitoQC compared to WT MEFs at the timepoints we examined. However, when BCL2L13 was deleted on top of the loss of the others, only then was a statistically significant loss of MitoQC-localized to lysosomes (**Fig. 4B**). Paralleling those results, only when all 4 mitophagy receptors were removed did we observe loss of mitolysosome formation by indirect immunofluorescence after CCCP. Furthermore, the residual mitophagy observed in the quadruple knockout suggests that other mitophagy receptors, such as FKBP8, PHB2^69–72^, or yet unidentified receptors, may contribute to CCCP-induced mitophagy.

Our findings also expand the contexts in which the BCL2L13 mitophagy receptor may be important, both to the many innate immune signaling contexts where TBK1 is critical (e.g. during pattern recognition receptor activation associated with infection, mtDNA release, etc.)^73,74^, as well as to conditions of low glucose and other metabolic stresses in which AMPK is activated. Similar to our previous findings with Ser108 of Parkin, we also observe here that Ser261 and Ser275 of BCL2L13 are being phosphorylated under conditions in which classic mitophagy is not well-established to be induced, such as after ETC complex inhibitors like rotenone or synthetic small molecule AMPK activators. It will be very interesting to examine whether phosphorylation of Ser261 alters recruitment of other mitophagy factors – perhaps even the ULK1 complex – to mitochondria. It is also worth noting that the two ten-amino-acid-long repeated sequences in front of the LIR in BCL2L13 contained many additional phosphorylation sites beyond Ser261, some of which like Ser259 were also robustly increased after CCCP. What the function of these repeat elements is will also be an interesting topic for future studies. What is clear is that a new model for the role of AMPK, TBK1, and ULK1 in triggering mitophagy has emerged from this and other recent studies (Fig. 4E). In response to hypoxia, ULK1 appears to play a prominent role and directly phosphorylates the serine in front of the LIR in BNIP3, NIX, and FUNCDC1, stimulating UB-independent mitophagy. In response to mitochondrial damage or synthetic AMPK activators, AMPK phosphorylates PARKIN and BCL2L13, stimulating both UB-dependent and independent mitophagy, while inhibiting ULK1-dependent activation of BNIP3, NIX, and FUNDC1. Finally multiple contexts, TBK1 can directly phosphorylate BCL2L13 to stimulate mitophagy. The interplay of these mitophagy receptors and their roles in human disease will continue to reveal critical insights in the coming years.

## Materials and Methods

### Antibodies and Reagents

BCL2L13 (16612-1-AP) was from ProteinTech. Cell Signaling Technology (CST) antibodies used were ACC (3662), phospho-ACC S79 (3661), AMPK (2532), phospho-AMPK T172 (2535), Raptor (2280), phospho-Raptor S792 (2083), TBK1 (3013), phospho-TBK1 S172 (5483), ULK1 (8054), phospho-ULK1 S757 (14202), phospho-ULK1 S555 (5869), ATG13 (13273), phospho-ATG13 Ser355 (46329), ATG14 (5504), phospho-ATG14 Ser29 (92340), VDAC (4661), TAX1BP1 (5105), NDP52 (60732), Optineurin (58981), phospho-Optineurin S177 (31304), FUNDC1 (49240), BNIP3 (3769), NIX (12396), HIF-1α (14179), ATG16L1 (8089), P62 (39749), and phospho-P62 S366 (26317). Phospho-BCL2L13 S261 and Phospho-BCL2L13 S275 were developed in collaboration with Gary Kasof at CST. Sigma-Aldrich antibodies used were beta-Actin (A5541) and Flag tag polyclonal (F7425). Phospho-ATG16L1-S278 was purchased from Abcam (ab195242). CCCP (C2759), rotenone (R8875). Compound 991 (GLXC-09267) was purchased from Glixx Laboratories Inc. Deferiprone (DFP) (S4067) was from Selleck Chemicals. MK-8722 (HY-111363) was from MedChem Express.

### Plasmids

The cDNA encoding human BCL2L13 was obtained from OriGene (RC200042). The Flag tag and attL1 sites (for BP Gateway reaction) were introduced by polymerase chain reaction (PCR) using standard methods. The resulting cDNAs were subcloned into pDONR221 with BP Clonase (Invitrogen, 11789020), and site-directed mutagenesis was performed using Q5 polymerase (New England Biolabs,M0491S). WT and mutant alleles in pDONR221 were sequenced in their entirety to verify no additional mutations were introduced during PCR or mutagenesis steps and then put into a Plenti destination vector (Addgene, 19067) by LR Gateway reaction (Invitrogen, 11791020 ). The pBabe.hygro-mCherry GFP-FIS1 (Mito-QC) construct was obtained from Professor Uri Manor at the Salk Institute, who originally obtained it from Professor Dario Alessi at the University of Dundee.

### Cell culture and cell lines

All cell lines were cultured in Dulbecco’s modified Eagle’s medium (DMEM) (Corning, MT10027CV) containing 10% fetal bovine serum (FBS) (Thermo Fisher Scientific, A5256701) and 1% penicillin/streptomycin (Thermo Fisher Scientific, 15140122) at 37°C in 5 % CO2. Small molecules were used at the indicated timepoints at the following concentrations: 991 (10uM), MK-8722 (10uM), rotenone (1uM), DFP (1mM), CCCP (20uM), and LPS (1ug/ml).

BCL2L13, NIX, BNIP3, FUNDC1, the quadruple knockout, and TBK1 knockout cells were generated using CRISPR-Cas9 editing with a ribonucleoprotein (RNP) complex. The RNP complex was composed of Cas9 Protein (Alt-R-Cas9, IDT), and a synthetic single-guide RNA (sgRNA) designed to target the gene of interest, and non-targeting guides were used for controls, which were synthesized by Synthego. Guides with high targeting scores and low probability of off-target effects were chosen. At least three independent sgRNA sequences were tested for each gene. The RNP complex was assembled by incubating Cas9 protein in a 1:1 molar ratio for 10 minutes at room temperature. The assembled RNP complex was then delivered into cells via electroporation. Single clones were cultured in 96 wells plates for 48-72 hours, followed by expansion into 6 well plates. Knockouts were then confirmed using immunoblotting. Successful clones were subsequently used for further assays.

### Stable expression of FLAG-tagged WT BCL2L13 and mutant BCL2L13

To generate stable cell lines expressing FLAG-tagged wild-type (WT) BCL2L13 and mutant BCL2L13, lentiviruses were produced by co-transfecting the lentiviral backbone constructs, along with the packaging plasmids pVSVg (Addgene, 8454) and pRev (Addgene, 12253), into HEK293T cells. Lipofectamine 2000 (ThermoFisher Scientific, 11668500) was used for transfection at a 3:1 lipofectamine/DNA ratio. Lentiviral supernatants were collected 72 hours post-transfection. MEF cells were infected with the filtered lentiviral supernatant (0.45 μm filter) supplemented with polybrene (5 µg/ml; Sigma-Aldrich, 107689) for 24 hours. After infection, cells were allowed to recover for 24 hours, followed by selection with neomycin (500 µg/ml; Invitrogen, 10131027). Stable cell lines were then validated for FLAG expression and levels of endogenous versus overexpressed BCL2L13 by Western blotting.

### Stable expression of mCherry-GFP-Fis1 (Mito-QC)

To stably express mCherry-GFP-Fis1 (mito-QC) in MEFs, retroviruses were generated by co-transfecting the pBabe mito-QC construct with the ampho packaging plasmid into HEK293T cells. The virus-containing supernatants were collected 48 hours post-transfection, filtered through 0.45 μm filters, and used to infect MEF cells in the presence of polybrene (5 µg/ml; Sigma-Aldrich, 107689). 48 hours post-infection, cells were sorted by flow cytometry (FACS;Aria Fusion), selecting cells that were positive for both GFP and mCherry, indicating successful expression of the Mito-QC construct.

### Western blotting

At experimental endpoints, whole cell lysates were generated by washing cells once with cold PBS. Cells were then harvested using cold CST lysis buffer (20mM Tris pH 7.5, 150mM NaCl, 1mM EDTA, 1mM EGTA, 1% Triton X-100, 2.5mM sodium pyrophosphate, 50mM NaF, 5mM β-glycero-phosphate, 50nM calyculin A, 1mM Na3VO4) supplemented with complete EDTA-free Protease Inhibitor Cocktail (Sigma-Aldrich, 11873580001) by cell scraping. Lysates were kept on ice and rotated at 4°C for 15 minutes. Following incubation, the lysates were centrifuged at 13,200 rpm at 4°C for 15 minutes to remove cell debris. The resulting supernatants were collected for BCA protein quantification and subsequent sample preparation for Western blot analysis.

### Mouse experiments for protein lysates from whole liver

All procedures using animals were approved by the Salk Institute Institutional Animal Care and Use Committee (IACUC) protocol 11-000029 to R.J.S. AMPKa1 and AMPKa2 floxed allele (Prkaa1^fl/fl^; Prkaa2^fl/fl^) or ULK1/2 floxed allele (Ulk1^fl/fl^; Ulk2^fl/fl^) mice bearing Ubc-creERT2 were treated with tamoxifen (3 mg/kg/day) or vehicle (control) for 5 consecutive days. Six weeks after tamoxifen injection, mice were fasted for 16 hours, refed for 1 hour, then for 3 hours with vehicle or MK-8722 (30 mg/kg) before being sacrificed as previously described^75^. Livers were collected, rapidly freeze-clamped, and lysates were prepared in lysis buffer for Western blot analysis.

### Immunoprecipitation

Cells stably expressing flag-tagged BCL2L13 were lysed in 2% CHAPS lysis buffer ((50 mM Tris-HCl, 50 mM NaCl, 1 mM EDTA, 2.0% CHAPS, 0.4 mM Na3VO4, 10 mM NaF, 10 mM sodium pyrophosphate, a protease inhibitor cocktail, pH 7.4), and samples were equilibrated. Flag-BCL2L13 was immunoprecipitated from 3 mg of lysates using Flag M2 magnetic beads (Sigma-Aldrich, M8823) at 4°C for 2h under rotation. Subsequently, the beads were washed three times with lysis buffer and two times with PBS. Protein complexes were eluted from beads using Laemmli sample buffer with 2% (v/v) beta-mercaptoethanol. Co-immunoprecipitating proteins were detected by immunoblot analysis.

### Mass Spectrometry

Flag BCL2L13 was overexpressed in HEK293T and U2OS cells, and immunoprecipitated with Flag magnetic beads (Sigma-Aldrich; M8823). Empty vector transfected cells were included as a negative control. Two quality controls were employed; 5% of the resulting IP samples were separated by SDS-PAGE gels, stained with coomassie blue (Abcam, ab119211), and another 5% was subjected to Western blotting to confirm the successful immunoprecipitation of flag-BCL2L13. The remaining IP samples were eluted in 2x SDS sample buffer and submitted to the Mass Spectrometry Core at Salk for targeted phospho-analysis to map phosphorylation sites on BCL2L13. Briefly, protein precipitate using methanol/chloroform protocol to remove unwanted materials from samples. Followed by resuspension of the protein pellet in urea and ammonium bicarb to reduce and alkylate, then digestion with GluC overnight, followed by reversed phase microcapillary/tandem mass spectrometry (LC/MS/MS). LC/MS/MS was performed using an EasynLC nanoflow HPLC (Proxeon Biosciences) with a self-packed 75 µm id x 15 cm C18 column coupled to a LTQ-Orbitrap XL mass spectrometer (Thermo Scientific) in the data-dependent acquisition and positive ion mode at 300 nL/min. Peptide ions from predicted phosphorylation sites were also targeted in MS/MS mode for quantitative analyses. MS/MS spectra collected via collision induced dissociation in the ion trap were searched against the concatenated target and decoy (reversed) single entry and full Swiss-Prot protein databases using Sequest (Proteomics Browser Software, Thermo Scientific) with differential modifications for Ser/Thr/Tyr phosphorylation (+79.97) and the sample processing artifacts Met oxidation (+15.99), deamidation of Asn and Gln (+0.984) and Cys alkylation (+57.02). Phosphorylated and un-phosphorylated peptide sequences were identified if they initially passed the following Sequest scoring thresholds against the target database: 1+7 ions, Xcorr ≥ 2.0 Sf ≥ 0.4, P ≥ 5; 2+ ions, Xcorr ≥ 2.0, Sf ≥ 0.4, P ≥ 5; 3+ ions, Xcorr ≥ 2.60, Sf ≥ 0.4, P ≥ 5 against the target protein database. Passing MS/MS spectra were manually inspected to ensure that all b- and y-fragment ions aligned with the assigned sequence and modification sites. Determination of the exact phosphorylation sites was aided using Fuzzy Ions and GraphMod and phosphorylation site maps were created using Protein Report software (Proteomics Browser Software suite, Thermo Scientific). False discovery rates (FDR) of peptide hits (phosphorylated and unphosphorylated) were estimated below 1.5% based on reversed database hits.

### Measurement of mitophagic flux by ratiometric flow cytometry

To assess the effect of knocking out mitophagy receptors on mitophagy, MEF cells stably expressing mCherry-GFP-Fis1 (to measure mitophagy based on degradation of the outer mitochondrial membrane) were analyzed using flow cytometry. Flow cytometry was performed with Aria fusion using the 488 nm and 561 nm lasers (Flow Cytometry Core, Salk). For each sample, 10,000 events were collected, and the appropriate side scatter (SSC) and forward scatter (FSC) profiles were used to exclude non-viable cells and doublets. Unstained cells were used to generate the ‘‘tandem+’’ gate. FlowJo software was then used to calculate the ratiometric analysis of mCherry/GFP within the ‘‘tandem+’’ gate. For quantification of mitochondrial delivery to lysosomes, the gate was set to 10% of tandem+ cells based on the mCherry/GFP ratio in vehicle-treated conditions.

### Immunofluorescence assay

At experimental endpoints, cells plated in 96-well imaging plates (Phenoplate, Revvity 6057302)were fixed with 4% paraformaldehyde (PFA) at room temperature (RT) for 15 min. Following fixation, wells were permeabilized with PBS + 0.1% Triton X-100 for 15 min at RT. Wells were blocked with 5% BSA and 10% goat serum in PBS (block buffer) for 30 min at RT. A cocktail of anti-HSP60 (EnCor Biotechnology, CPCA-HSP60, 1:500), and anti-RAB7 (Cell Signaling Technology, E907E, 1:500) primary antibodies was prepared in block buffer and incubated in each well for 1 hour at RT. Wells were washed with PBS and incubated with secondary antibodies, anti-Chicken IgY AF647 (Abcam, ab150171, 1:500), anti-Rabbit IgG AF488 (Invitrogen, A11008, 1:500) and 4’,6-diamidino-2-phenylindole (DAPI) counterstain (Thermo Fisher Scientific, 62248, 1:1000) in block buffer at RT for 30 min. Wells were washed with PBS and imaged within one week of sample generation.

### Automated high-content confocal microscopy

MEFs were plated in 96-well imaging plates (Phenoplate, Revvity 6057302) and stained according to the immunofluorescence assay protocol described above. Fixed cells were imaged with an Olympus 60X air interface objective on a BioTek Cytation C10 Confocal Imaging Reader (Agilent) automated spinning disk confocal microscope equipped with a Hamamatsu ORCA CMOS digital camera. Single optical sections were imaged for each acquisition field, with consistent focus maintained by laser autofocus. 16 acquisition fields were randomly selected per well. Representative images have been deconvolved using 5-iteration default deconvolution in Gen5 software. Representative images are scaled evenly regarding brightness and contrast between experimental groups.

### Quantitative microscopy analysis of mitolysosomes

Confocal micrographs were quantified using open-source image analysis software Cellprofiler^76^. The Cellprofiler pipeline associated with this manuscript is available in Supplementary Materials (File S1). Image analysis was performed on raw confocal micrographs. Single-cell analysis was performed by segmentation of nuclear objects based on DAPI intensity, using the “IdentifyPrimaryObjects” module, followed by propagation of the nuclear objects to the cell border based on a low-stringency threshold of the HSP60 stain in the “IdentifySecondaryObjects” module. Next, the mitochondrial network was segmented based on adaptive (Otsu) thresholding of the HSP60 immunostain using the “IdentifyPrimaryObjects” module. The “MeasureObjectSizeShape” module was then used to define the size distribution of mitochondrial objects, and objects with a size of <1 μm^2^ were designated as “mitochondrial fragments” using the “FilterObjects” module. Using the MeasureObjectIntensity, the mean fluorescence intensity (MFI) of the RAB7 immunostain was recorded. The “FilterObjects” module was used to select RAB7^+^ mitochondrial fragments (MFI > 3 standard deviations above the mean DMSO-treated control group), which were designated as “mitolysosomes.” Following segmentation and filtering of mitolysosomes, features were summarized to cell-level means and related to the single cell using the “RelateObjects” module.

## Acknowledgements

We thank Elijah Trefts, Chien-Min Hung, and Kristina Hellberg from the Shaw lab for the generation of lysates from WT, AMPK-, and ULK1/2-liver KO mice used in Fig. 2. We particularly thank Alan Tung, who pioneered this project as a UCSD undergraduate and generated several reagents to initiate this work. We would also like to thank Professor Christina Towers and Anvita Komarla for their assistance with the mito-QC and flow cytometry analysis, and Uri Manor for providing the mito-QC construct. This study was supported by grants to R.J.S. from the National Institutes of Health (NIH) R35CA220538 and P01CA120964 and grant to G.S.S RO1 AG077324. M.B.R. was supported by the Hewitt Foundation Postdoctoral Fellowship. This work was also supported by the Waitt Advanced Biophotonics Core Facility of the Salk Institute with funding from NCI CCSG: P30 CA014195 and the Waitt Foundation, the Flow Cytometry Core Facility of the Salk Institute with funding from NCI CCSG P30 CA01495, the Mass Spectrometry Core Facility of the Salk Institute with funding from NCI CCSG P30 CA01495 and instrumentation resources at the UC San Diego Agilent Center of Excellence in Cellular Intelligence.

## Abbreviations

CCCP: Carbonyl Cyanide m-Chlorophenyl Hydrazone;
LPS: Lipopolysaccharide;
DFP: Deferiprone;
OMM: Outer mitochondrial membrane;
ULK1: Unc-51 Like Autophagy Activating Kinase 1;
AMPK: AMP-activated protein kinase;
TBK1: TANK-binding kinase 1;
PINK1: PTEN-induced kinase 1;
BNIP3L: BCL2 Interacting Protein 3 Like (NIX);
BNIP3: BCL2 Interacting Protein 3;
FUNDC1: Fun14 Domain Containing 1;
BCL2L13: BCL2-like 13;
LIR: LC3 interacting region;
MEF: Mouse embryonic fibroblast;
ETC: Electron Transport Chain;
HEK293: Human embryonic kidney;
SLR: Sequestosome/p62-like receptors.

---

**Figure S1.**
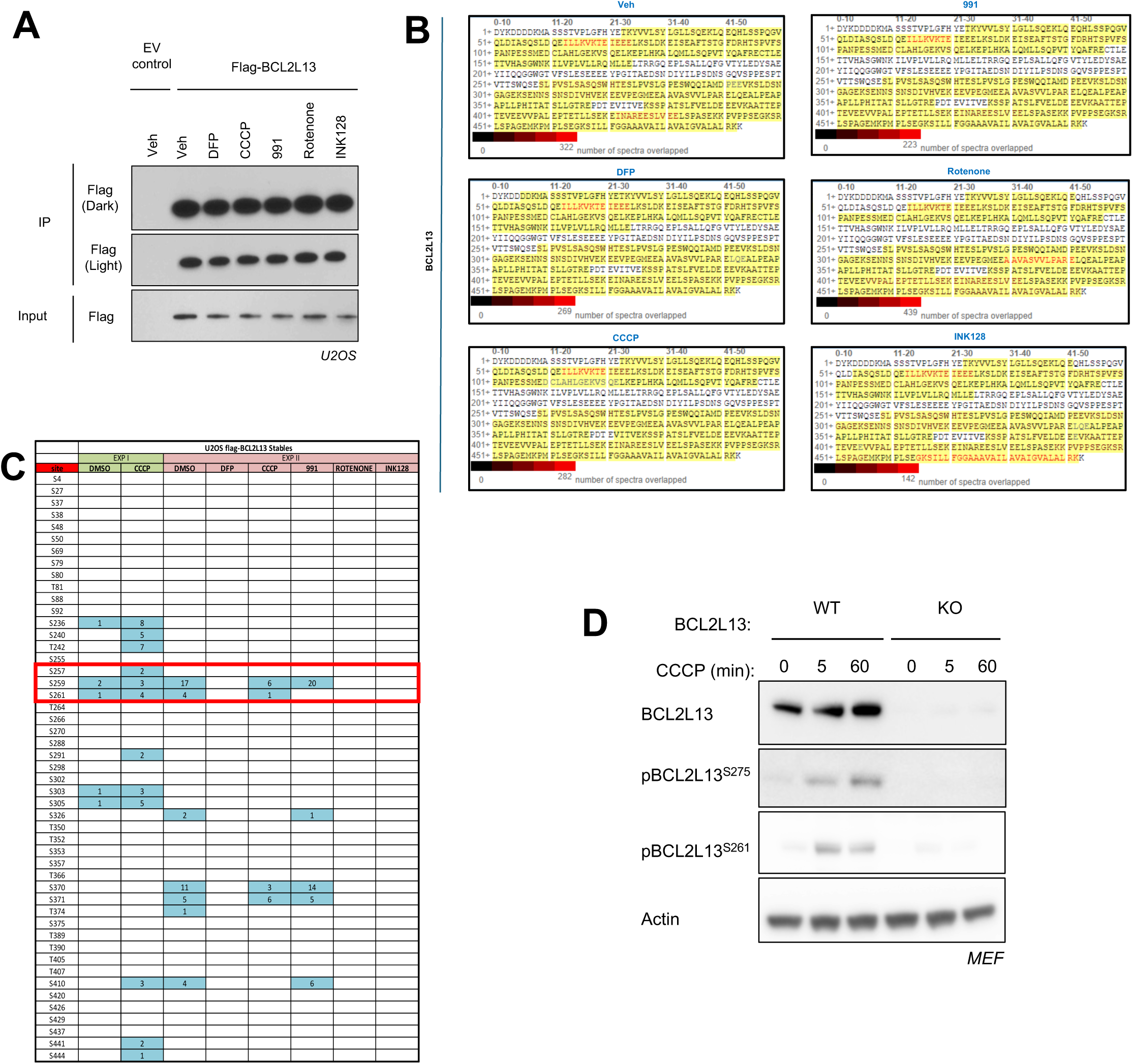
Mass Spectrometry Identification of CCCP-Induced Phosphorylation Sites on BCL2L13. **(A)** Immunoblot of U2OS cells stably expressing flag-tagged BCL2L13, treated with various autophagy and mitophagy-inducing compounds: 20 µM CCCP, 1 mM DFP, 10 µM 991, 100 ng/ml rotenone, and 1 µM INK128 for 1 hour. After treatment, cells were lysed in 2% CHAPS buffer, and the lysates were subjected to flag immunoprecipitation (IP). **(B)** Protein sequence coverage map of the BCL2L13 samples submitted for mass spectrometry analysis. GluC digestion resulted in 75% coverage of the BCL2L13 protein sequence. **(C)** The table summarizes the spectral counts for each detected phosphorylation site. Notably, three phosphorylation sites — S275, S259, and S261 — show a significant increase in spec count after CCCP treatment compared to DMSO vehicle. These sites match the consensus kinase motifs for ULK1- or TBK1-like kinases, as highlighted in the red box. **(D)** Immunoblots of MEF cells from wild-type (WT) or CRISPR-Cas9 RNP-mediated BCL2L13 knockout (KO) lines, treated with 20 µM CCCP at the indicated time-point to validate the phospho-specific antibody developed by CST. These blots demonstrate that the antibody specifically detects BCL2L13 at serine 275 (S275) in MEF cells, with phosphorylation levels increasing following treatment with CCCP.

**Figure S2.**
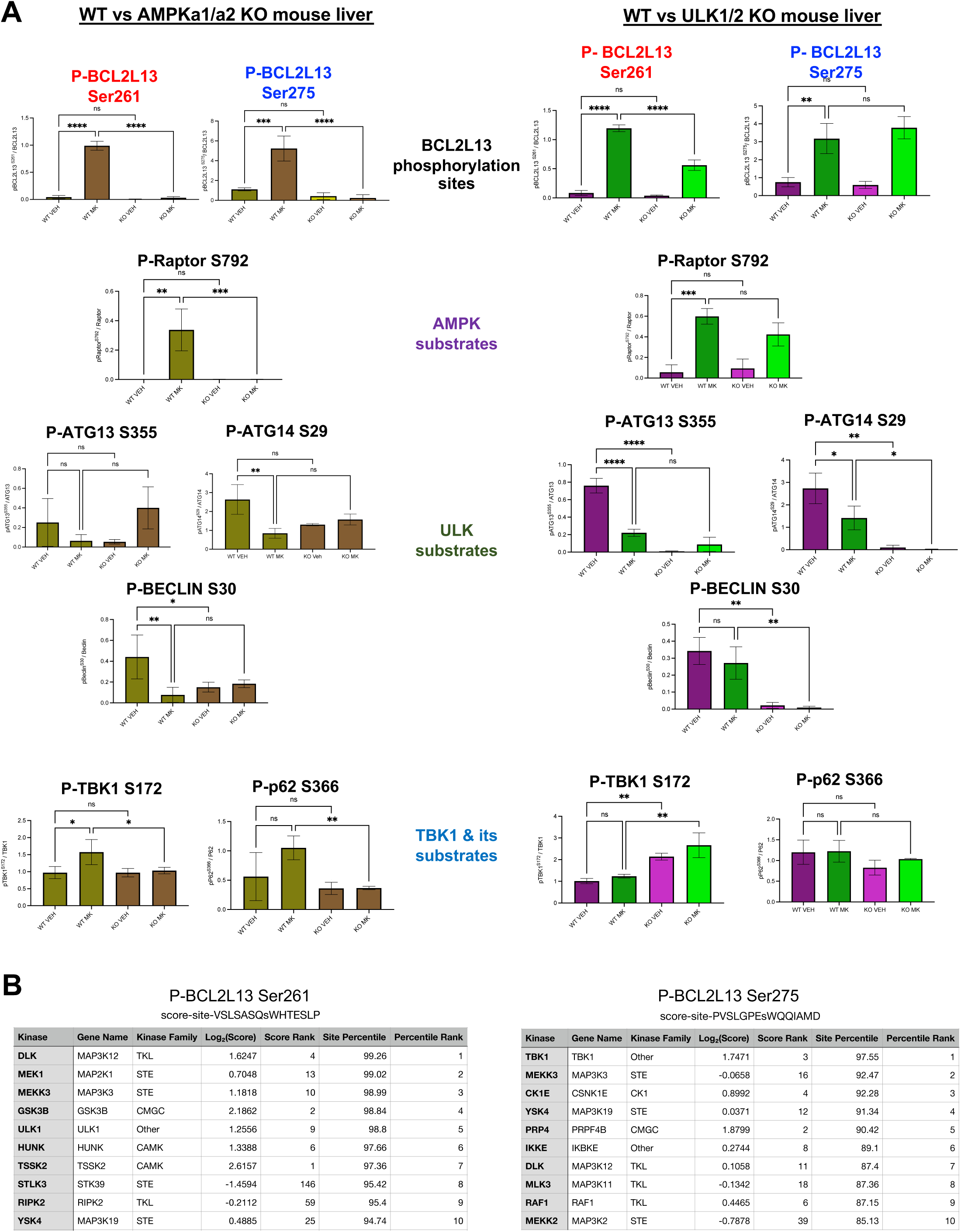
Quantification of BCL2L13 phosphorylation and predicted kinases for Bcl2L13 Ser261 and Ser275. **(A)** AMPK, ULK and TBK1 signaling in mouse livers of indicated genotypes following treatment of mice with vehicle or the AMPK director activator MK-877 (one dose, 30mg/kg). This analysis corresponds to the Western blot data shown in Densitometric analysis was performed to assess protein expression levels. Phospho-protein levels were quantified and normalized to total protein levels and loading control, Actin. Data are presented as mean ± SEM. Statistical analysis was performed using one-way ANOVA. Asterisks indicate statistically significant differences compared to the control group (\**p* < 0.05, ** p < 0.01, *** p < 0.001, **** p< 0.0001). **(B)** Kinase library (ref) top 10 predicted kinases for Ser261 and Ser275

**Figure S3.**
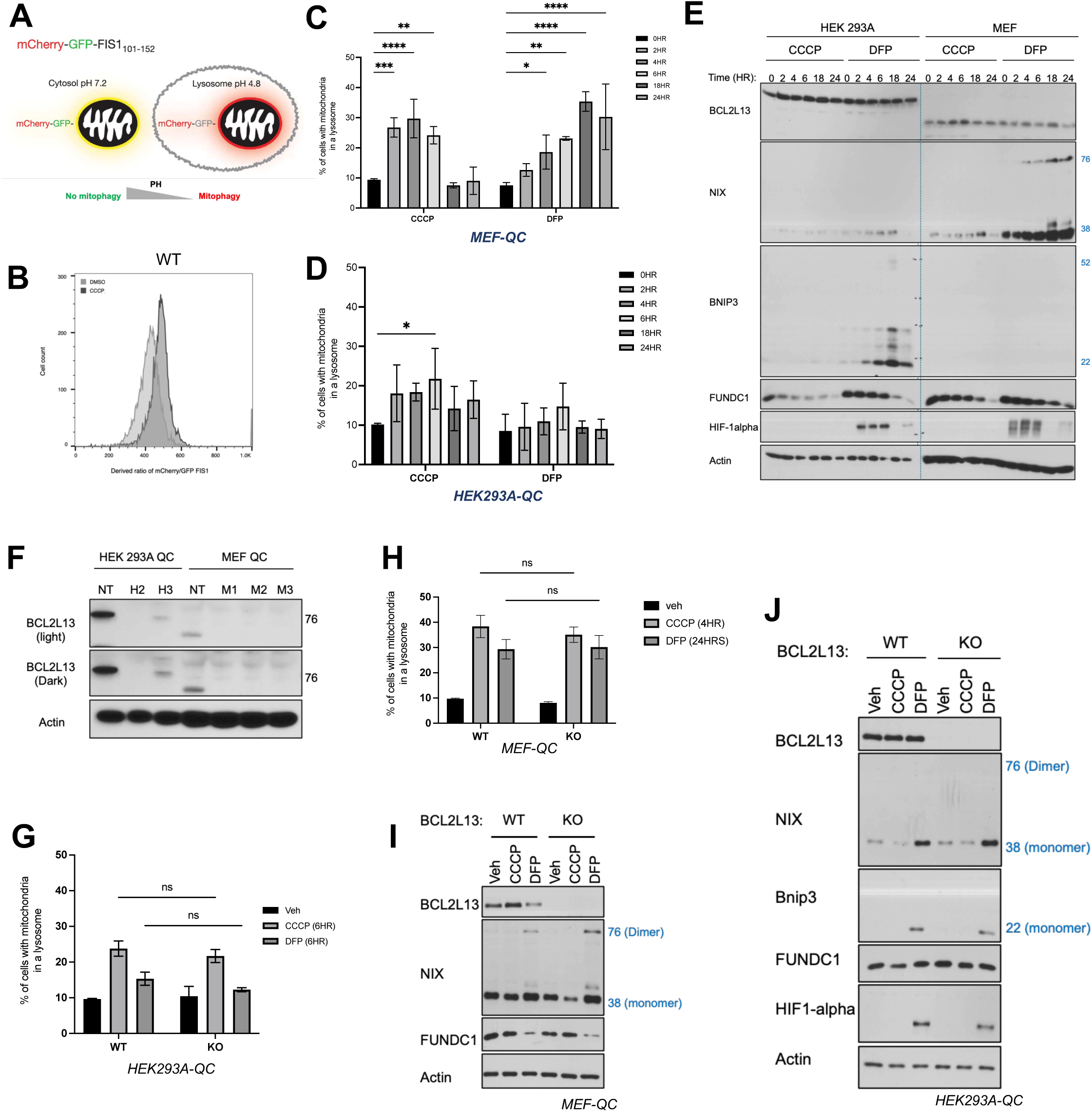
Mito-QC-based mitophagy assay in MEF and HEK293A cells. (**A**) Diagram of the Mito-QC construct: mCherry-GFP tandem mitophagy assay. (**B)** Representative histogram of ratiometric flow cytometry analysis before and after 4 hours of CCCP in WT MEF cells stably expressing the mitophagy reporter Mito-QC (“MEF-QC”). **(C)** MEF-QC cells were treated with CCCP or DFP at the indicated time points. Following treatment, cells were harvested in flow buffer, and mitophagy was quantified using flow cytometry based on the mCherry/GFP ratio. (**D**) HEK293A-QC cells treated and analyzed as in panel C. **(E)** Immunoblot of protein levels of the mitophagy receptors in MEF and HEK293A following CCCP or DFP at the indicated timepoints. (**F)** Human and mouse BCL2L13 CRISPR-Cas9 deletion was assessed in MEF-QC and HEK293A-QC cells. **(G)** WT or CRISPR-Cas9 mediated BCL2L13 knockout (KO) HEK293-QC cells were treated with CCCP or DFP at the indicated time points. Following treatment, cells were harvested in flow buffer, and mitophagy was quantified using flow cytometry based on the mCherry/GFP ratio. (**H)** WT or BCL2L13 KO MEF-QC cells were treated with CCCP or DFP at the indicated time points and mitophagy was quantified using flow cytometry based on the mCherry/GFP ratio as in panel G. **(I)** Immunoblot analysis of mitophagy receptors in WT and BCL2L13 KO MEFs reveals differential regulation of ubiquitin-independent mitophagy receptors MEF cells exhibit differential regulation of the ubiquitin-independent mitophagy receptors. **(J)** Immunoblot analysis of mitophagy receptors in WT and BCL2L13 KO HEK293A cells. Ratiometric flow cytometry data (mCherry/GFP ratio) are presented as the mean ± SEM for biological replicates (N = 3). Statistical significance was assessed using Two-Way ANOVA. (\**p* < 0.05, ** p < 0.01, *** p < 0.001, **** p< 0.0001).

**Figure S4.**
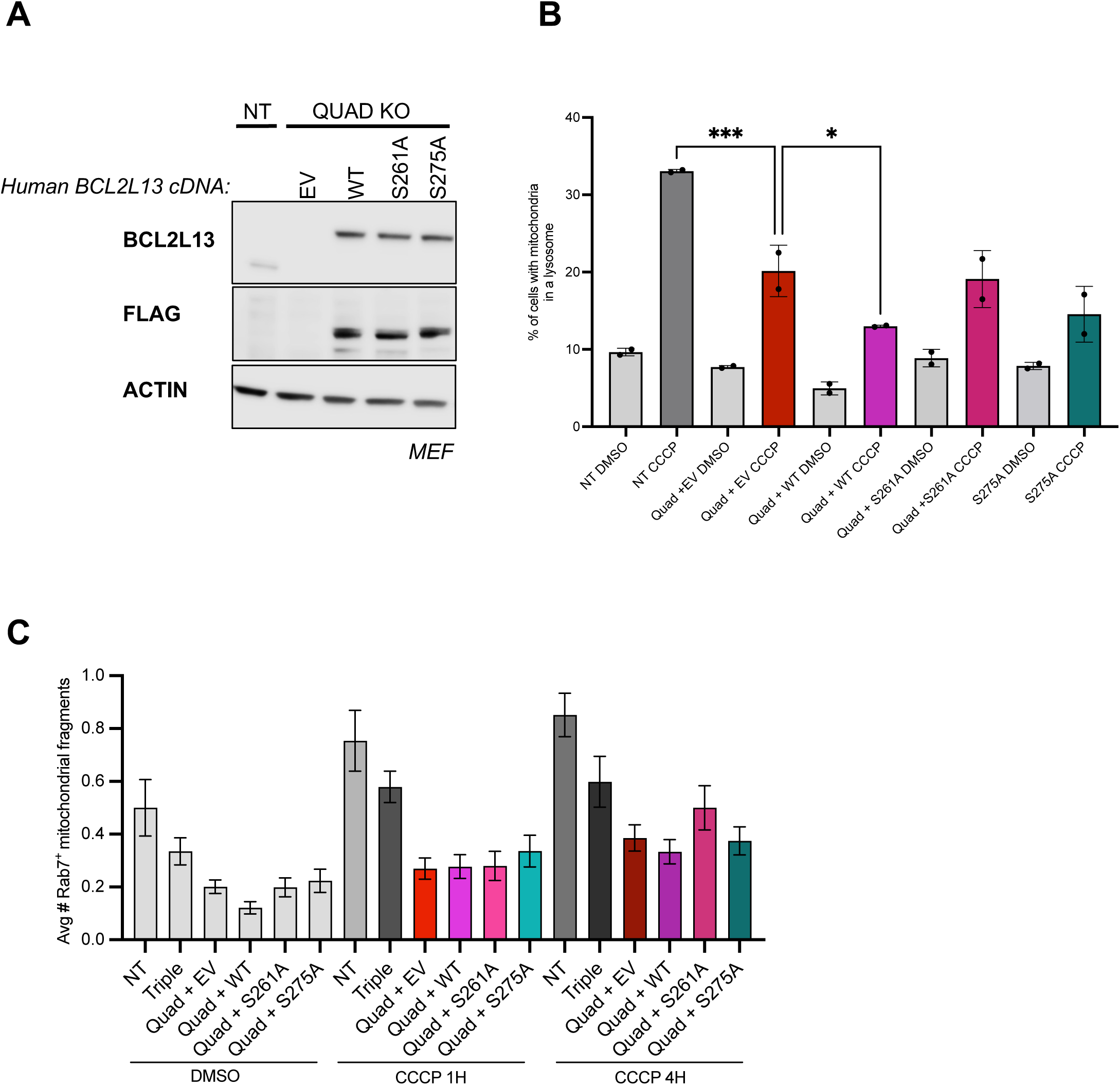
Evaluation of mitophagy in BCL2L13 wild-type and Ser-Ala mutant addback in the quadruple KO MEFs. Using MitoQC flow cytometry analysis and quantitative co-locatization of HSP60 and RAB7 by high-throughput indirect immunofluorescence imaging, as in Fig. 4. **(A)** Endogenous mouse BCL2L13 levels (NT cells) alongside stable expression of WT human Flag-tagged BCL2L13 and Serine-to-Alanine mutants in the Quad KO MEFs. Endogenous BCL2L13 protein levels are comparable to the flag-tagged addbacks. Note that human BCL2L13 runs ∼20kD larger than mouse BCL2L13. **(B)** Mitophagy in Quad KO MEFs stably expressing the mitophagy reporter (mito-QC) with addback of BCL2L13 WT point mutants was analyzed using flow cytometry. Quantitative analysis of ratiometric flow cytometry (mCherry/GFP). Data are presented as mean ± SEM. Statistical analysis was performed using one-way ANOVA. Asterisks indicate statistically significant differences compared to the control group (\**p* < 0.05, ** p < 0.01, *** p < 0.001, **** p< 0.0001). **(C)** Control (NT), Triple KO, and Quad KO MEF stably expressing with indicated BCL2L13 cDNAs were treated with 20 µM CCCP for 1H or 4H timepoints and immunostained for HSP60 and Rab7, as detailed in Fig. 4. Mitolysosome formation was quantified as RAB7+ mitochondrial fragments using CellProfiler. Bar graphs represent the mean and standard error of >200 cells/condition.

## Notes

### Competing Interest Statement

The authors have declared no competing interest.

## REFERENCES

1. Um J-H, Yun and J. Emerging role of mitophagy in human diseases and physiology. BMB Rep 2017; 50:299–307.

2. Gustafsson ÅB, Dorn GW. Evolving and Expanding the Roles of Mitophagy as a Homeostatic and Pathogenic Process. Physiol Rev 2019; 99:853–92.

3. Onishi M, Yamano K, Sato M, Matsuda N, Okamoto K. Molecular mechanisms and physiological functions of mitophagy. Embo J 2021; 40:e104705.

4. Choubey V, Zeb A, Kaasik A. Molecular Mechanisms and Regulation of Mammalian Mitophagy. Cells 2021; 11:38.

5. Pickles S, Vigié P, Youle RJ. Mitophagy and Quality Control Mechanisms in Mitochondrial Maintenance. Curr Biol 2018; 28:R170–85.

6. Zachari M, Ktistakis NT. Mammalian Mitophagosome Formation: A Focus on the Early Signals and Steps. Frontiers Cell Dev Biology 2020; 8:171.

7. Kubli DA, Gustafsson ÅB. Mitochondria and Mitophagy. Circ Res 2012; 111:1208–21.

8. Ney PA. Mitochondrial autophagy: Origins, significance, and role of BNIP3 and NIX. Biochim Biophys Acta (BBA) - Mol Cell Res 2015; 1853:2775–83.

9. Ktistakis NT. The dynamics of mitochondrial autophagy at the initiation stage. Biochem Soc T 2021; 49:2199–210.

10. Montava-Garriga L, Ganley IG. Outstanding Questions in Mitophagy: What We Do and Do Not Know. J Mol Biol 2020; 432:206–30.

11. Wanderoy S, Hees JT, Klesse R, Edlich F, Harbauer AB. Kill one or kill the many: interplay between mitophagy and apoptosis. Biol Chem 2020; 402:73–88.

12. Montava-Garriga L, Ganley IG. Outstanding Questions in Mitophagy: What We Do and Do Not Know. J Mol Biol 2020; 432:206–30.

13. Belousov DM, Mikhaylenko EV, Somasundaram SG, Kirkland CE, Aliev G. The Dawn of Mitophagy: What Do We Know by Now? Curr Neuropharmacol 2020; 19:170–92.

14. Heo J-M, Ordureau A, Paulo JA, Rinehart J, Harper JW. The PINK1-PARKIN Mitochondrial Ubiquitylation Pathway Drives a Program of OPTN/NDP52 Recruitment and TBK1 Activation to Promote Mitophagy. Mol Cell 2015; 60:7–20.

15. Fritsch LE, Moore ME, Sarraf SA, Pickrell AM. Ubiquitin and Receptor-Dependent Mitophagy Pathways and Their Implication in Neurodegeneration. J Mol Biol 2020; 432:2510–24.

16. Hung C-M, Lombardo PS, Malik N, Brun SN, Hellberg K, Nostrand JLV, Garcia D, Baumgart J, Diffenderfer K, Asara JM, et al. AMPK/ULK1-mediated phosphorylation of Parkin ACT domain mediates an early step in mitophagy. Sci Adv 2021; 7:eabg4544.

17. Longo M, Bishnu A, Risiglione P, Montava-Garriga L, Cuenco J, Sakamoto K, MacKintosh C, Ganley IG. Opposing roles for AMPK in regulating distinct mitophagy pathways. Mol Cell 2024; 84:4350–4367.e9.

18. Marinković M, Novak I. A brief overview of BNIP3L/NIX receptor-mediated mitophagy. Febs Open Bio 2021; 11:3230–6.

19. McWilliams TG, Prescott AR, Montava-Garriga L, Ball G, Singh F, Barini E, Muqit MMK, Brooks SP, Ganley IG. Basal Mitophagy Occurs Independently of PINK1 in Mouse Tissues of High Metabolic Demand. Cell Metab 2018; 27:439–449.e5.

20. Lampert MA, Orogo AM, Najor RH, Hammerling BC, Leon LJ, Wang BJ, Kim T, Sussman MA, Gustafsson ÅB. BNIP3L/NIX and FUNDC1-mediated mitophagy is required for mitochondrial network remodeling during cardiac progenitor cell differentiation. Autophagy 2019; 15:1182–98.

21. Chen M, Chen Z, Wang Y, Tan Z, Zhu C, Li Y, Han Z, Chen L, Gao R, Liu L, et al. Mitophagy receptor FUNDC1 regulates mitochondrial dynamics and mitophagy. Autophagy 2016; 12:689–702.

22. Novak I, Kirkin V, McEwan DG, Zhang J, Wild P, Rozenknop A, Rogov V, Löhr F, Popovic D, Occhipinti A, et al. Nix is a selective autophagy receptor for mitochondrial clearance. Embo Rep 2010; 11:45–51.

23. Li M, Jia J, Zhang X, Dai H. Selective binding of mitophagy receptor protein Bcl-rambo to LC3/GABARAP family proteins. Biochem Biophys Res Commun 2020; 530:292–300.

24. Rogov VV, Nezis IP, Tsapras P, Zhang H, Dagdas Y, Noda NN, Nakatogawa H, Wirth M, Mouilleron S, McEwan DG, et al. Atg8 family proteins, LIR/AIM motifs and other interaction modes. Autophagy Rep 2023; 2:2188523.

25. Rogov V, Dötsch V, Johansen T, Kirkin V. Interactions between Autophagy Receptors and Ubiquitin-like Proteins Form the Molecular Basis for Selective Autophagy. Mol Cell 2014; 53:167–78.

26. Zhu Y, Massen S, Terenzio M, Lang V, Chen-Lindner S, Eils R, Novak I, Dikic I, Hamacher-Brady A, Brady NR. Modulation of Serines 17 and 24 in the LC3-interacting Region of Bnip3 Determines Pro-survival Mitophagy versus Apoptosis*. J Biol Chem 2013; 288:1099–113.

27. Wu W, Tian W, Hu Z, Chen G, Huang L, Li W, Zhang X, Xue P, Zhou C, Liu L, et al. ULK1 translocates to mitochondria and phosphorylates FUNDC1 to regulate mitophagy. Embo Rep 2014; 15:566–75.

28. Rogov VV, Suzuki H, Marinković M, Lang V, Kato R, Kawasaki M, Buljubašić M, Šprung M, Rogova N, Wakatsuki S, et al. Phosphorylation of the mitochondrial autophagy receptor Nix enhances its interaction with LC3 proteins. Sci Rep 2017; 7:1131.

29. Matsumoto G, Wada K, Okuno M, Kurosawa M, Nukina N. Serine 403 Phosphorylation of p62/SQSTM1 Regulates Selective Autophagic Clearance of Ubiquitinated Proteins. Mol Cell 2011; 44:279–89.

30. Pilli M, Arko-Mensah J, Ponpuak M, Roberts E, Master S, Mandell MA, Dupont N, Ornatowski W, Jiang S, Bradfute SB, et al. TBK-1 Promotes Autophagy-Mediated Antimicrobial Defense by Controlling Autophagosome Maturation. Immunity 2012; 37:223–34.

31. Hu L, Xie H, Liu X, Potjewyd F, James LI, Wilkerson EM, Herring LE, Xie L, Chen X, Cabrera JC, et al. TBK1 Is a Synthetic Lethal Target in Cancer with VHL Loss. Cancer Discov 2020; 10:460–75.

32. Moore AS, Holzbaur ELF. Dynamic recruitment and activation of ALS-associated TBK1 with its target optineurin are required for efficient mitophagy. Proc Natl Acad Sci 2016; 113:E3349–58.

33. Richter B, Sliter DA, Herhaus L, Stolz A, Wang C, Beli P, Zaffagnini G, Wild P, Martens S, Wagner SA, et al. Phosphorylation of OPTN by TBK1 enhances its binding to Ub chains and promotes selective autophagy of damaged mitochondria. Proc Natl Acad Sci 2016; 113:4039–44.

34. Vargas JNS, Hamasaki M, Kawabata T, Youle RJ, Yoshimori T. The mechanisms and roles of selective autophagy in mammals. Nat Rev Mol Cell Biol 2023; 24:167–85.

35. Lamark T, Johansen T. Mechanisms of Selective Autophagy. Annu Rev Cell Dev Biol 2021; 37:1– 27.

36. Goodall EA, Kraus F, Harper JW. Mechanisms underlying ubiquitin-driven selective mitochondrial and bacterial autophagy. Mol Cell 2022; 82:1501–13.

37. Ganley IG, Simonsen A. Diversity of mitophagy pathways at a glance. J Cell Sci 2022; 135:jcs259748.

38. Poole LP, Bock-Hughes A, Berardi DE, Macleod KF. ULK1 promotes mitophagy via phosphorylation and stabilization of BNIP3. Sci Rep-uk 2021; 11:20526.

39. Kataoka T, Holler N, Micheau O, Martinon F, Tinel A, Hofmann K, Tschopp J. Bcl-rambo, a Novel Bcl-2 Homologue That Induces Apoptosis via Its Unique C-terminal Extension*. J Biol Chem 2001; 276:19548–54.

40. Warren CFA, Wong-Brown MW, Bowden NA. BCL-2 family isoforms in apoptosis and cancer. Cell Death Dis 2019; 10:177.

41. Kataoka T. Biological properties of the BCL-2 family protein BCL-RAMBO, which regulates apoptosis, mitochondrial fragmentation, and mitophagy. Frontiers Cell Dev Biology 2022; 10:1065702.

42. Terešak P, Lapao A, Subic N, Boya P, Elazar Z, Simonsen A. Regulation of PRKN-independent mitophagy. Autophagy 2022; 18:24–39.

43. Meng F, Sun N, Liu D, Jia J, Xiao J, Dai H. BCL2L13: physiological and pathological meanings. Cell Mol Life Sci 2021; 78:2419–28.

44. Otsu K, Murakawa T, Yamaguchi O. BCL2L13 is a mammalian homolog of the yeast mitophagy receptor Atg32. Autophagy 2015; 11:1932–3.

45. Murakawa T, Okamoto K, Omiya S, Taneike M, Yamaguchi O, Otsu K. A Mammalian Mitophagy Receptor, Bcl2-L-13, Recruits the ULK1 Complex to Induce Mitophagy. Cell Reports 2019; 26:338–345.e6.

46. Bellot G, Garcia-Medina R, Gounon P, Chiche J, Roux D, Pouysségur J, Mazure NM. Hypoxia-Induced Autophagy Is Mediated through Hypoxia-Inducible Factor Induction of BNIP3 and BNIP3L via Their BH3 Domains. Mol Cell Biol 2009; 29:2570–81.

47. Madhu V, Boneski PK, Silagi E, Qiu Y, Kurland I, Guntur AR, Shapiro IM, Risbud MV. Hypoxic Regulation of Mitochondrial Metabolism and Mitophagy in Nucleus Pulposus Cells Is Dependent on HIF-1α–BNIP3 Axis. J Bone Miner Res 2020; 35:1504–24.

48. Wu W, Li W, Chen H, Jiang L, Zhu R, Feng D. FUNDC1 is a novel mitochondrial-associated-membrane (MAM) protein required for hypoxia-induced mitochondrial fission and mitophagy. Autophagy 2016; 12:1675–6.

49. Johansen T, Lamark T. Selective Autophagy: ATG8 Family Proteins, LIR Motifs and Cargo Receptors. J Mol Biol 2020; 432:80–103.

50. Allen GFG, Toth R, James J, Ganley IG. Loss of iron triggers PINK1/Parkin-independent mitophagy. Embo Rep 2013; 14:1127–35.

51. Egan DF, Chun MGH, Vamos M, Zou H, Rong J, Miller CJ, Lou HJ, Raveendra-Panickar D, Yang C-C, Sheffler DJ, et al. Small Molecule Inhibition of the Autophagy Kinase ULK1 and Identification of ULK1 Substrates. Mol Cell 2015; 59:285–97.

52. Helgason E, Phung QT, Dueber EC. Recent insights into the complexity of Tank-binding kinase 1 signaling networks: The emerging role of cellular localization in the activation and substrate specificity of TBK1. FEBS Lett 2013; 587:1230–7.

53. Gwinn DM, Shackelford DB, Egan DF, Mihaylova MM, Mery A, Vasquez DS, Turk BE, Shaw RJ. AMPK Phosphorylation of Raptor Mediates a Metabolic Checkpoint. Mol Cell 2008; 30:214–26.

54. Herzig S, Shaw RJ. AMPK: guardian of metabolism and mitochondrial homeostasis. Nat Rev Mol Cell Biol 2018; 19:121–35.

55. Garcia D, Shaw RJ. AMPK: Mechanisms of Cellular Energy Sensing and Restoration of Metabolic Balance. Mol Cell 2017; 66:789–800.

56. Garcia D, Hellberg K, Chaix A, Wallace M, Herzig S, Badur MG, Lin T, Shokhirev MN, Pinto AFM, Ross DS, et al. Genetic Liver-Specific AMPK Activation Protects against Diet-Induced Obesity and NAFLD. Cell Rep 2019; 26:192–208.e6.

57. Durand JK, Zhang Q, Baldwin AS. Roles for the IKK-Related Kinases TBK1 and IKKε in Cancer. Cells 2018; 7:139.

58. Herhaus L. TBK1 (TANK-binding kinase 1)-mediated regulation of autophagy in health and disease. Matrix Biol 2021; 100:84–98.

59. Runde AP, Mack R, S.J. PB, Zhang J. The role of TBK1 in cancer pathogenesis and anticancer immunity. J Exp Clin Cancer Res 2022; 41:135.

60. Zhang Q, Liu S, Zhang C-S, Wu Q, Yu X, Zhou R, Meng F, Wang A, Zhang F, Chen S, et al. AMPK directly phosphorylates TBK1 to integrate glucose sensing into innate immunity. Mol Cell 2022; 82:4519–4536.e7.

61. Vargas JNS, Wang C, Bunker E, Hao L, Maric D, Schiavo G, Randow F, Youle RJ. Spatiotemporal Control of ULK1 Activation by NDP52 and TBK1 during Selective Autophagy. Mol Cell 2019; 74:347–362.e6.

62. Wild P, Farhan H, McEwan DG, Wagner S, Rogov VV, Brady NR, Richter B, Korac J, Waidmann O, Choudhary C, et al. Phosphorylation of the Autophagy Receptor Optineurin Restricts Salmonella Growth. Science 2011; 333:228–33.

63. Goodwin JM, Dowdle WE, DeJesus R, Wang Z, Bergman P, Kobylarz M, Lindeman A, Xavier RJ, McAllister G, Nyfeler B, et al. Autophagy-Independent Lysosomal Targeting Regulated by ULK1/2-FIP200 and ATG9. Cell Rep 2017; 20:2341–56.

64. Murakawa T, Ito J, Rusu M-C, Taneike M, Omiya S, Moncayo-Arlandi J, Nakanishi C, Sugihara R, Nishida H, Mine K, et al. AMPK regulates Bcl2-L-13-mediated mitophagy induction for cardioprotection. Cell Rep 2024; 43:115001.

65. McWhirter SM, Fitzgerald KA, Rosains J, Rowe DC, Golenbock DT, Maniatis T. IFN-regulatory factor 3-dependent gene expression is defective in Tbk1-deficient mouse embryonic fibroblasts. Proc Natl Acad Sci 2004; 101:233–8.

66. Murakawa T, Yamaguchi O, Hashimoto A, Hikoso S, Takeda T, Oka T, Yasui H, Ueda H, Akazawa Y, Nakayama H, et al. Bcl-2-like protein 13 is a mammalian Atg32 homologue that mediates mitophagy and mitochondrial fragmentation. Nat Commun 2015; 6:7527.

67. Fujiwara M, Tian L, Le PT, DeMambro VE, Becker KA, Rosen CJ, Guntur AR. The mitophagy receptor Bcl-2–like protein 13 stimulates adipogenesis by regulating mitochondrial oxidative phosphorylation and apoptosis in mice. J Biol Chem 2019; 294:12683–94.

68. Johnson JL, Yaron TM, Huntsman EM, Kerelsky A, Song J, Regev A, Lin T-Y, Liberatore K, Cizin DM, Cohen BM, et al. An atlas of substrate specificities for the human serine/threonine kinome. Nature 2023; 613:759–66.

69. Aguilera MO, Robledo E, Melani M, Wappner P, Colombo MI. FKBP8 is a novel molecule that participates in the regulation of the autophagic pathway. Biochim Biophys Acta (BBA) - Mol Cell Res 2022; 1869:119212.

70. Bhujabal Z, Birgisdottir ÅB, Sjøttem E, Brenne HB, Øvervatn A, Habisov S, Kirkin V, Lamark T, Johansen T. FKBP8 recruits LC3A to mediate Parkin-independent mitophagy. EMBO Rep 2017; 18:947–61.

71. Yoo S, Yamashita S, Kim H, Na D, Lee H, Kim SJ, Cho D, Kanki T, Jung Y. FKBP8 LIRL-dependent mitochondrial fragmentation facilitates mitophagy under stress conditions. FASEB J 2020; 34:2944–57.

72. Wei Y, Chiang W-C, Sumpter R, Mishra P, Levine B. Prohibitin 2 Is an Inner Mitochondrial Membrane Mitophagy Receptor. Cell 2017; 168:224–238.e10.

73. Perry AK, Chow EK, Goodnough JB, Yeh W-C, Cheng G. Differential Requirement for TANK-binding Kinase-1 in Type I Interferon Responses to Toll-like Receptor Activation and Viral Infection. J Exp Med 2004; 199:1651–8.

74. Liu S, Cai X, Wu J, Cong Q, Chen X, Li T, Du F, Ren J, Wu Y-T, Grishin NV, et al. Phosphorylation of innate immune adaptor proteins MAVS, STING, and TRIF induces IRF3 activation. Science 2015; 347:aaa2630.

75. Hardie DG, Schaffer BE, Brunet A. AMPK: An Energy-Sensing Pathway with Multiple Inputs and Outputs. Trends Cell Biol 2016; 26:190–201.

76. Lamprecht MR, Sabatini DM, Carpenter AE. CellProfiler: free, versatile software for automated biological image analysis. BioTechniques 2007; 42:71–5.

